# Therapeutic targeting of PGBD5-induced DNA repair dependency in pediatric solid tumors

**DOI:** 10.1101/181040

**Authors:** Anton G. Henssen, Casie Reed, Eileen Jiang, Heathcliff Dorado Garcia, Jennifer von Stebut, Ian C. MacArthur, Patrick Hundsdoerfer, Jun Hyun Kim, Elisa de Stanchina, Yasumichi Kuwahara, Hajime Hosoi, Neil Ganem, Filemon Dela Cruz, Andrew L. Kung, Johannes H. Schulte, John H. Petrini, Alex Kentsis

**Author notes:** Correspondence: Alex Kentsis, MD, PhD.

## Abstract

Despite intense efforts, the cure rates of childhood and adult solid tumors are not satisfactory. Resistance to intensive chemotherapy is common, and targets for molecular therapies are largely undefined. We have now found that the majority of childhood solid tumors, including rhabdoid tumors, neuroblastoma, medulloblastoma and Ewing sarcoma, express an active DNA transposase *PGBD5* that can promote site-specific genomic rearrangements in human cells. Using functional genetic approaches, we found that mouse and human cells deficient in non-homologous end joining (NHEJ) DNA repair cannot tolerate the expression of PGBD5. In a chemical screen of DNA damage signaling inhibitors, we identified AZD6738 as a specific sensitizer of PGBD5-dependent DNA damage and apoptosis. We found that expression of PGBD5, but not its nuclease activity-deficient mutant, was sufficient to induce hypersensitivity to AZD6738. Depletion of endogenous PGBD5 conferred resistance to AZD6738 in human tumor cells. PGBD5-expressing tumor cells accumulated unrepaired DNA damage in response to AZD6738 treatment, and underwent apoptosis in both dividing and G1 phase cells in the absence of immediate DNA replication stress. Accordingly, AZD6738 exhibited nanomolar potency against the majority of neuroblastoma, medulloblastoma, Ewing sarcoma and rhabdoid tumor cells tested, while sparing non-transformed human and mouse embryonic fibroblasts *in vitro*. Finally, treatment with AZD6738 induced apoptosis and regression of human neuroblastoma and medulloblastoma tumors engrafted in immunodeficient mice *in vivo*. This effect was potentiated by combined treatment with cisplatin, including significant anti-tumor activity against patient-derived primary neuroblastoma xenografts. These findings delineate a therapeutically actionable synthetic dependency induced in PGBD5-expressing solid tumors.

## Introduction

In spite of the improvements in intensive combination chemotherapy, surgery and radiotherapy, the treatment of the majority of childhood and adult solid tumors remains inadequate. For example, neuroblastomas and medulloblastomas characterized by amplifications of the *MYCN* and *MYC* oncogenes, respectively, remain mostly fatal (*1-3*). Likewise, cancers defined by mutations of the genes encoding the SWI/SNF chromatin remodeling complex, such as rhabdoid tumors, are almost uniformly incurable (*4*). Finally, the majority of human sarcomas, if they cannot be removed completely by surgery, such as Ewing sarcoma for example, tend to be chemotherapy resistant and lethal (*5*). The majority of refractory childhood solid tumors are characterized by mutations of factors that regulate gene expression or complex genomic rearrangements, both of which are not generally amenable to current pharmacologic strategies. Thus, new therapeutic approaches are urgently needed to improve the cure rates for these patients.

To enhance the current therapeutic index, synthetic cellular relationships have been leveraged for cancer therapy (*6*). For example, tumors with inefficient homologous recombination DNA repair due to mutations of *BRCA1/2* exhibit synthetic lethality with inhibitors of poly ADP-ribose polymerases (PARPs), enabling significant improvements in the treatment of patients as a result of clinical PARP inhibitors (*7, 8*). In addition, synthetic dependencies in metabolic function (*9*), chromatin remodeling (*10*), and DNA damage signaling (*11-13*), are beginning to be explored to develop improved targeted therapies. In particular, intrinsic DNA damage due to oncogene or replication stress such as MYC (*14*), and tumorigenic deficiencies in the DNA damage response due to mutations of *TP53*, *ATM* or *ATR* have been found to confer susceptibility to specific inhibitors of DNA damage repair signaling (*15*). However, these mutations are generally rare in pediatric cancers, and little is known about therapeutically targetable synthetic dependencies in childhood solid tumors.

Recently, the human *piggyBac transposable element derived 5* (*PGBD5*) was identified as an active DNA transposase that is able to mobilize synthetic DNA transposons in human cells (*16*). PGBD5-mediated DNA transposition requires the DDD catalytic triad in the PGBD5 transposase domain and specific DNA recognition sequences and target sites (*16*). In particular, PGBD5 was found to be expressed in the majority of childhood solid tumors, including refractory rhabdoid tumors, where it promotes site-specific genomic rearrangements and mutations of tumor suppressor genes at least in part due to the aberrant targeting of its DNA nuclease activity (*17*). This tumorigenic nuclease activity of PGBD5 raises the possibility that PGBD5-expressing cells may depend on active DNA damage repair and signaling.

Here, we report that PGBD5 activity confers a functional dependence on the KU complex that binds DNA double-strand breaks (DSBs), and the ATR and ATM kinases that control DNA damage repair signaling in cells. We found that PGBD5 activity is sufficient to confer this synthetic dependence, and endogenous PGBD5 expression is necessary to render childhood solid tumor cells susceptible to inhibitors of DSB repair signaling. As a result, its pharmacologic targeting using selective inhibitors of DNA damage signaling exhibits therapeutic activity in multiple preclinical models of neuroblastoma, medulloblastoma, Ewing sarcoma, and rhabdoid tumors that express PGBD5 *in vitro* and *in vivo*. The availability of clinical-grade inhibitors of DNA damage signaling offers immediate potential for translation into clinical trials for patients with refractory childhood solid tumors, the majority of which express PGBD5, as well as distinct subsets of PGBD5-expressing adult solid cancers.

## Results

### PGBD5-expressing cells do not tolerate deficiency of non-homologous end-joining DNA repair

Eukaryotic DNA transposases rely on cellular DNA repair mechanisms to restore intact target sites upon DNA rearrangements (*18*). In mammalian cells, this activity is principally carried out by the classic non-homologous end joining (NHEJ) DSB repair apparatus (*19*). NHEJ repair consists of the heterodimeric KU70/KU80 complex that binds DSB ends, and the DNA damage repair signaling factors including the ataxia telangiectasia mutated (ATM) and ATM-and Rad3-related (ATR) kinases that in concert lead to chromatin reorganization and DNA ligase-mediated DSB repair (*20*). Depending on the specific molecular features of DNA damage and cell state, assembly of different repair complexes can lead to activation of specific signaling pathways (*21-23*). This suggests that distinct forms of intrinsic DNA damage in cancer cells can be used selectively for their synthetic lethal targeting.

To test the cellular DNA repair requirements of PGBD5, we used mouse embryonic fibroblasts (MEFs) from mice deficient for *Ku80*^*-/-*^, *Atm*^*-/-*^, and hypomorphic Seckel syndrome allele of *Atr*^*S/S*^, and immortalized with the SV40 large T antigen (*24-26*) (Fig. 1A). Similar experiments using *Lig4*^*-/-*^ MEFs could not be performed because of their severe proliferation defect (data not shown) (*27*). We used a doxycycline-inducible transgene encoding human *PGBD5*, and confirmed equal PGBD5 protein expression upon doxycycline induction using Western immunoblotting (Fig. 1A). Wild-type SV40 large T antigen-immortalized MEFs exhibited no measureable changes in cell growth upon doxycycline-induced PGBD5 expression (Figs. 1B-E). In contrast, *Atm*^*-/-*^, *Atr*^*S/S*^, and *Ku80*^*-/-*^ MEFs underwent cell death, as detected by the significant accumulation of cleaved caspase 3 (*p* = 1.0e-2, 8.0e-3, and 1.0e-3, respectively, Figs. 1B-C), terminal deoxynucleotidyl transferase dUTP nick end labeling (TUNEL; *p* = 1.0e-3 and 2.0e-3, respectively, Fig. 1D), and histone H2AX S139 phosphorylation (γH2AX; *p* = 3.0e-3 and 2.0e-2, respectively, Fig. 1E). Deficiency of *Ku80* that functions in direct DSB binding during NHEJ DNA repair exhibited similar levels of cell death as the respective deficiencies of *Atm* and *Atr* that contribute to the activation and propagation of DNA damage signaling (Fig. 1B). Thus, PGBD5 expression requires the cellular NHEJ and DNA damage signaling apparatus.

**Figure 1.**
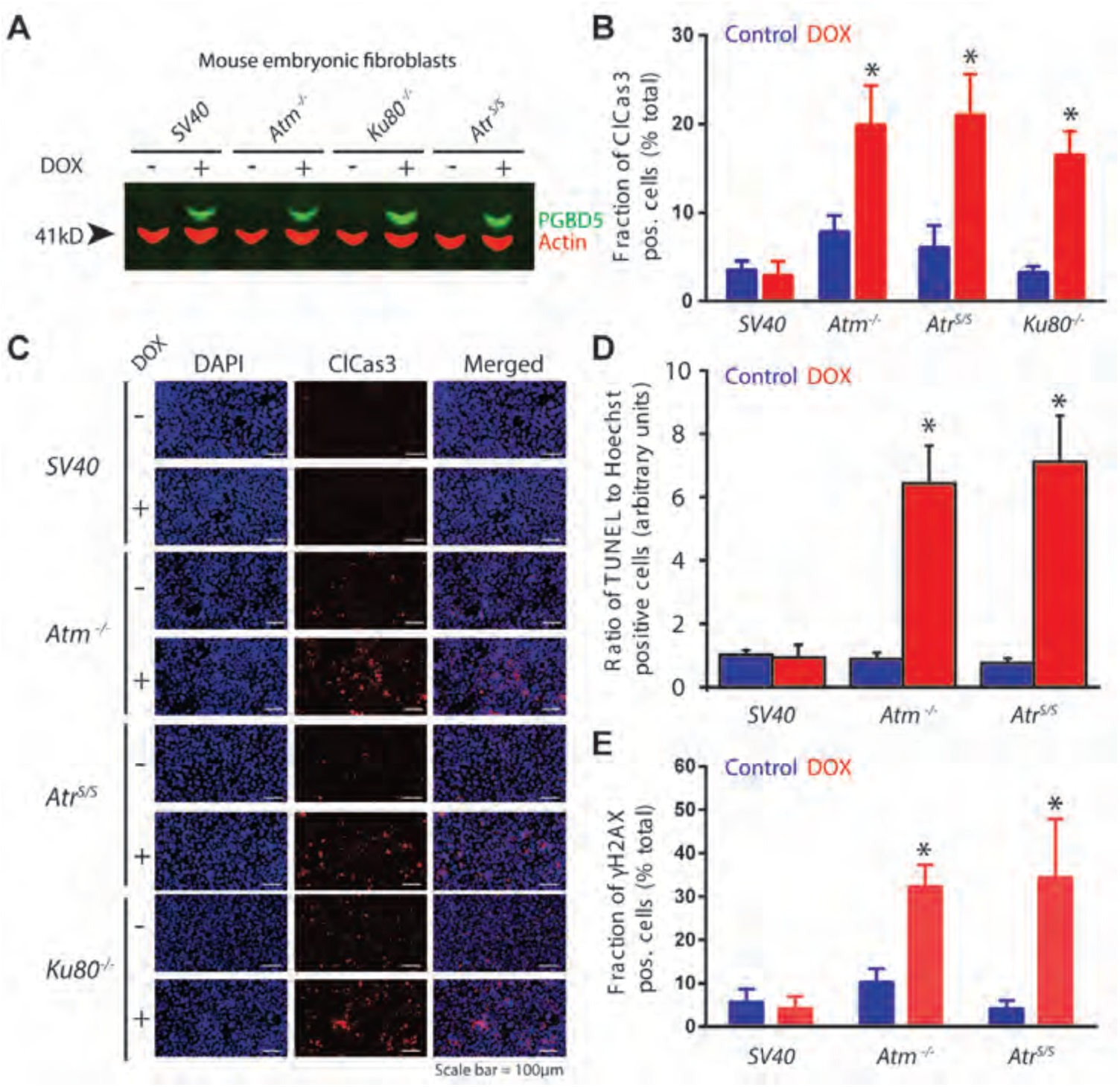
PGBD5-expressing cells do not tolerate deficiency of non-homologous end-joining DNA repair. **(A)** Western blot of PGBD5 protein expression after induction with doxycycline (500 ng/ml for 24 hours) of SV40 large T antigen-immortalized mouse embryonic fibroblasts deficient for *Atm*^*-/-*^, *Atr*^*S/S*^ or *Ku80*^*-/-*^. Actin serves as loading control. **(B)** Induction of apoptosis as measured by cleaved caspase-3 staining of mouse embryonic fibroblasts deficient DNA repair factors upon doxycycline-induced (48 hours) expression of PGBD5 (red) as compared to control PBS-treated cells (blue). \**p* = 0.010, 0.008, and 0.0010 for *Atm*^*-/-*^, *Atr*^*S/S*^*, and Ku80*^*-/-*^ of doxycycline vs. control, respectively. **(C)** Representative photomicrographs of mouse embryonic fibroblasts upon doxycycline-induced PGBD5 expression for 48 hours (+) as compared to PBS-treated controls (-), as stained for DAPI (blue) and cleaved caspase-3 (red). Scale bar = 100 μm. **(D-E)** Induction of DNA DSB as measured by TUNEL staining (D) and γH2AX (E) of mouse embryonic fibroblasts deficient DNA repair factors upon doxycycline-induced (48 hours) expression of PGBD5 (red) as compared to control PBS-treated cells (blue). \**p* = 0.0010 and 0.0020 for *Atm*^*-/-*^ and *Atr*^*S/S*^ of doxycycline vs. control, respectively (D). \**p* = 0.0030 and 0.020 for *Atm*^*-/-*^ and *Atr*^*S/S*^ of doxycycline vs. control, respectively (E). Error bars represent standard deviations of three biologic replicates.

### PGBD5 expression is sufficient to confer susceptibility to inhibition of DNA damage repair signaling

Based on the finding that cells expressing PDBD5 are dependent on active NHEJ DNA damage repair and signaling, we hypothesized that PGBD5 expression might render cells susceptible to pharmacological inhibition of DNA damage signaling. To test this idea, we used primary human retinal pigment epithelial (RPE) cells immortalized with telomerase that allow the investigation of DNA damage response in genomically stable, non-transformed human cells (*28*). We reasoned that specific PGBD5-dependent DNA damage response requirements may be identified by comparative analysis of growth and survival of RPE cells expressing *GFP-PGBD5*, as compared to *GFP* control and its catalytically inactive *GFP-PGBD5* D168A/D194A/D386A mutant. This mutant is expressed equally to wild-type PGBD5, as assessed by Western immunoblotting, and physically associates with chromatin, as assessed by chromatin immunoprecipitation and DNA sequencing (ChIP-seq), but does not support DNA transposition in reporter assays (*17*). To minimize the possible contribution of secondary effects of PGBD5 expression due to its induction of mutagenic DNA rearrangements and cell transformation, we used cells for experiments immediately after transgene transduction (*17*).

Thus, we screened a panel of commercially available inhibitors of DNA damage signaling in their ability to selectively interfere with the growth and survival of RPE cells expressing *GFP-PGBD5* as compared to wild-type or control transgene expressing cells (Fig. 2A & S1, Table S1). In particular, we observed that the ATR-and ATM-selective kinase inhibitors AZD6738 and KU60019 exhibited more than 20-fold and 5-fold enhanced activity, respectively, against RPE cells expressing *GFP-PGBD5* as compared *GFP* control (Fig. 2A). Consistent with the notion that tolerance of PGBD5 DNA nuclease activity requires active DNA repair in cells, mutation of the aspartate catalytic triad (D168A, D194A, D386A) thought to catalyze phosphodiester bond hydrolysis during transposase-induced DNA rearrangements (*16*), completely abrogated the enhanced susceptibility of RPE cells upon PGBD5 expression (Fig. 2B). We confirmed that the activity of AZD6738 is at least in part due to its selective inhibition of ATR as compared to ATM (*29, 30*), insofar as *Atm*^*-/-*^ and *Atr*^*S/S*^ MEFs exhibited respectively increased and diminished susceptibilities to AZD6738, in agreement with the known relationship between ATM and ATR kinase signaling (*11-13*) (*IC*_*50*_ = 0.023 and 2.0 μM, respectively, as compared to 0.36 μM for wild-type control; Fig. 2C). Thus, PGBD5-dependent effects may be explained by the selective inhibition of DNA damage signaling by ATR-and ATM-selective AZD6738 and KU60019, respectively. Lack of PGBD5-dependent effects of other albeit potent DNA damage signaling inhibitors is presumably related to their respective selectivity profiles, such as for example AZ20 which potently inhibits ATM, ATR and mTOR, and VE-822 (VX-970) which inhibits both ATR and ATM kinases (Table S1).

**Figure 2.**
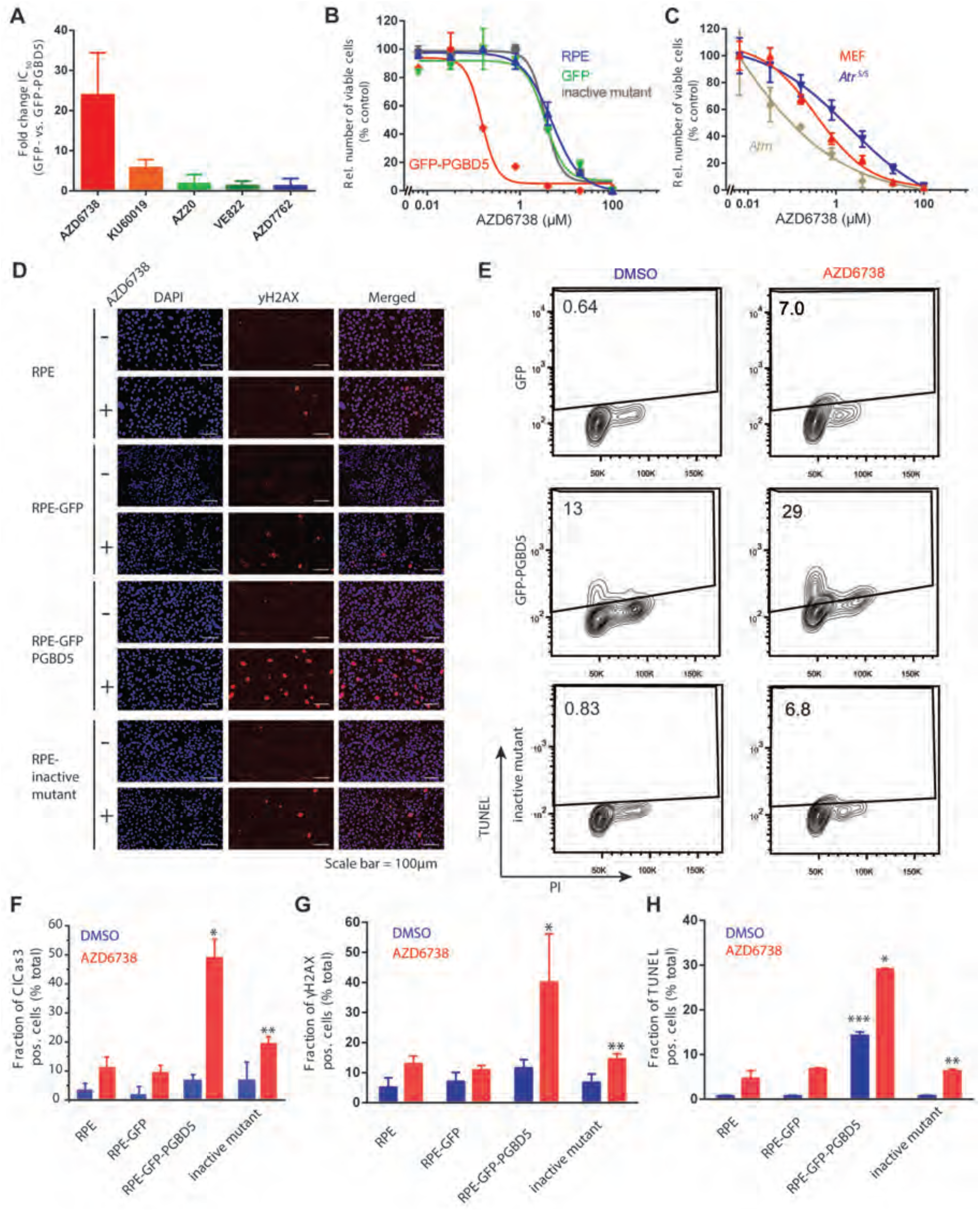
PGBD5 expression is sufficient to confer sensitivity to DNA damage signaling inhibition. **(A)** Ratios of 50% inhibitory concentrations (*IC*_*50*_) upon 120 hours of drug treatment with the DNA damage signaling inhibitors AZD6738, KU60019, AZ20, VE822, and AZD7762 of RPE cells expressing *GFP-PGBD5* or *GFP* control. Ratios of 1 indicate equal susceptibility of *GFP-PGBD5* as compared to control *GFP*-expressing cells. **(B)** Dose response cell viability curves of RPE cells (blue) expressing *GFP-PGBD5* (red) as compared to *GFP* control (green) or catalytically inactive mutant (D168A/D194A/D386A, black) *GFP-PGBD5* treated with AZD6738 for 120 hours.**(C)** Dose response cell viability curves of wild-type MEFs immortalized with SV40 large T antigen (red), as compared to *Atm*^*-/-*^ (brown) and *Atr*^*S/S*^ (blue) treated with AZD6738 for 120 hours. **(D)** Representative photomicrographs of RPE cells upon treatment with 500 nM AZD6738 for 72 hours (+) as compared to DMSO-treated controls (-), and expression of *GFP*, *GFP-PGBD5*, or inactive mutant (D168A/D194A/D386A) *GFP-PGBD5*, as stained for DAPI (blue) and γH2AX (red). Scale bar = 100 μm. **(E)** Flow cytometric analysis of TUNEL and propidium iodide staining of RPE cells expression *GFP-PGBD5* as compared to control *GFP* or inactive mutant (D168A/D194A/D386A) *GFP-PGBD5*, and treated with 500 nM AZD6738, as compared to DMSO control for 48 hours. Percentages of TUNEL-positive cells are labeled as indicated. **(F-H)** Induction of apoptosis and DNA damage, as measured by cleaved caspase 3 (F), γH2AX (G), and TUNEL staining (H) of RPE cells treated with 500 nM AZD6738 for 48 hours (red), as compared to DMSO control (blue). Expression of *GFP-PGBD5* as compared to control *GFP* leads to significant induction of caspase 3 cleavage, γH2AX, and TUNEL \**p* = 0.00030, 0.040, and 5.6 x 10^-6^, respectively. Mutation of the catalytic D168A/D194A/D386A (inactive mutant) *GFP-PGBD5* rescues this effect \*\**p* = 0.030, 0.010, and 1.5 x 10^-5^, respectively. Expression of *GFP-PGBD5* but not its inactive mutant or control *GFP* causes significant increase in TUNEL staining in the absence of AZD6738 treatment \*\*\**p* = 1.2 x 10^-6^. Error bars represent standard deviations of three biologic replicates.

Importantly, as with the functional genetic requirements of DNA repair and DNA damage signaling induced by PGBD5 expression in MEFs (Fig. 1), we found that active DNA damage signaling blocked by AZD6738 is also required for the growth and survival of PGBD5-expressing human RPE cells (Figs. 2D-H). Specifically, we found that RPE cells expressing *GFP-PGBD5*, but not those expressing *GFP* or inactive mutant *GFP-PGBD5*, exhibited significantly increased DNA damage upon treatment with AZD6738, as measured γH2AX staining analysis (*p* = 0.040, Fig. 2D & G). Similarly, we observed significantly increased TUNEL labeling in RPE cells expressing *GFP-PGBD5* upon treatment with AZD6738, whereas cells expressing control *GFP* or the catalytically inactive *GFP-PGBD5* mutant exhibited unperturbed steady-state background TUNEL levels (*p* = 5.6 x 10^-6^, Fig. 2E & H). Consistent with the notion that wild-type PGBD5 is actively inducing DSBs in cells, we observed significantly increased TUNEL levels even in the absence of AZD6738 treatment, an effect that was completely abolished by the mutation of its putative nuclease catalytic triad (Figs. 2E & H). Likewise, TUNEL accumulation upon PGBD5 expression and AZD6738 treatment was predominantly observed in G1 phase of the cell cycle (Fig. 2E). In accordance with the accumulation of unrepaired DNA damage upon AZD6738 treatment in PGBD5-expressing cells, we observed significantly increased levels of apoptosis, as measured by caspase 3 cleavage, in cells expressing *GFP-PGBD5*, as compared to those expressing its inactive mutant or *GFP* control (*p* = 3.0e-4, Fig. 2F). Consistent with the lack of apparent effect of AZD6738 on TUNEL levels in S phase cells (Fig. 2E), we confirmed that the PGBD5-specific susceptibility to AZD6738 was not immediately associated with DNA replication stress, as assessed by Western immunoblotting of phosphorylated RPA32 T21 and S4/8, as compared to the DNA topoisomerase I inhibitor camptothecin that predominantly induces DSBs during DNA replication in S phase (*31*) (Fig. 3). In agreement with this, RPE cells expressing *GFP-PGBD5* did not show increased cell cycling as compared to cells expressing *GFP* control, as measured by 5-ethynyl-2’-deoxyuridine (EdU) incorporation (Fig. S2). Lastly, in agreement with site-specific accumulation of DSBs (*32*), we observed mostly punctate in contrast to pan-nuclear γH2AX accumulation in PGBD5-expressing cells treated with AZD6738 (Fig. S3). Thus, PGBD5 expression is sufficient to confer specific susceptibility to pharmacologic inhibitors of DNA damage signaling such as AZD6738.

**Figure 3.**
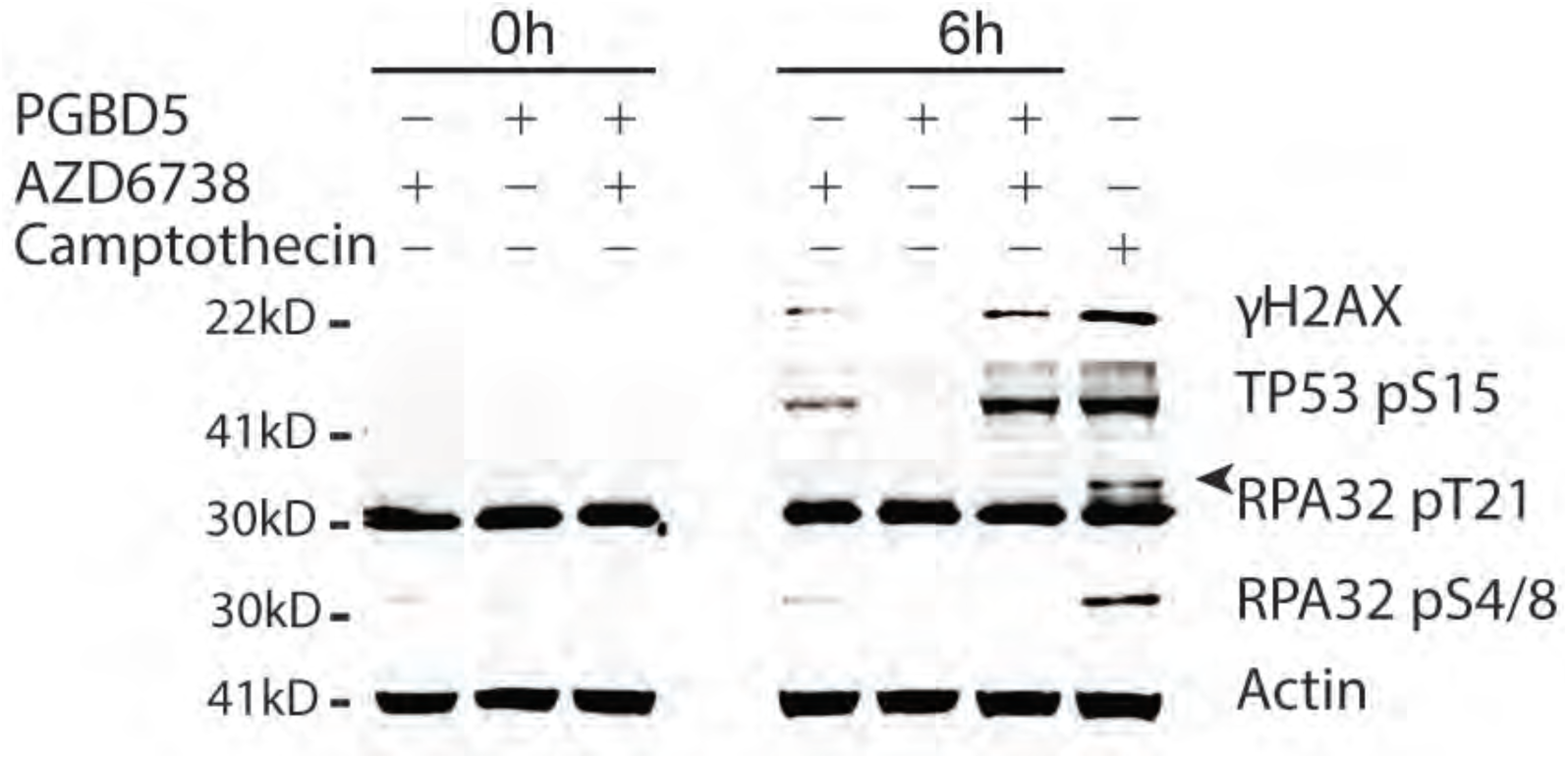
Treatment with AZD6738 induces PGBD5-dependent DNA damage. Western immunoblot of RPE cells expressing *GFP-PGBD5* or *GFP* and treated with 500 nM AZD6738 for 6 hours, as indicated. PGBD5 expression increases levels of γH2AX and TP53 pS15 upon combination with AZD6738 treatment, but not RPA32 pT21 or pS4/8. Actin serves as loading control, and treatment with 1.5 μM camptothecin for 2 hours serves as positive control for replication stress-mediated induction of RPA32 phosphorylation. Arrow head marks the specific RPA32 pT21 band.

### Rhabdoid tumor, medulloblastoma, neuroblastoma, and Ewing sarcoma cells that express *PGBD5* exhibit enhanced sensitivity to AZD6738

Previously, we observed that PGBD5 is expressed in the majority of childhood solid tumors, including neuroblastoma, medulloblastoma, Ewing sarcoma, and rhabdoid tumors (*17*). In particular, in rhabdoid tumors, we found that the DNA recombinase activity of PGBD5 was necessary and sufficient to induce genomic rearrangements in both rhabdoid tumor cell lines and patient tumors. We have now found that expression of PGBD5 in non-transformed mouse embryonic fibroblasts and human RPE cells is sufficient to induce DNA damage as measured by TUNEL incorporation in cells, which can be potentiated by DNA damage signaling inhibitor AZD6738. Thus, we reasoned that AZD6738 may exhibit anti-tumor activity against childhood solid tumor cells expressing PGBD5.

To test this idea, we treated a panel of rhabdoid tumor, neuroblastoma, medulloblastoma and Ewing sarcoma cell lines, as well as non-transformed human RPE and BJ cells with AZD6738. We observed that 19 childhood tumor cell lines tested exhibited enhanced sensitivity to AZD6738, with 50% inhibitory concentrations (*IC*_*50*_) largely in the nanomolar range (Fig. 4A & S4, Table S2). In contrast, non-transformed RPE and BJ cells were relatively resistant to AZD6738, with *IC*_*50*_ values in the high micromolar range (Fig. 4A & S4). Indeed, the susceptibility to AZD6738, as measured by its *IC*_*50*_ values *in vitro*, exhibited a significant correlation with the level of PGBD5 protein expression, as assessed by quantitative fluorescent Western immunoblotting (*p* = 4.4e-3, Fig. 4B, S4, S5). This association did not appear to segregate with tumor tissue type (Fig. 4B), or the presence of mutations in genes known to affect DNA damage signaling, such as *TP53*, *ATM*, *ATR, MYC, MYCN, XRCC3, XRCC5, CHK1, BRCA2, RAS,* and *ATRX* (Fig. S5 and Table S2). In addition, since ATM deficiency can confer increased sensitivity to AZD6738 (*12*), we confirmed that PGBD5 expression did not affect the expression of ATM itself in RPE cells, and that human tumor cells exhibiting enhanced susceptibility to AZD6738 lacked *ATM* mutations and retained ATM protein expression (Fig. S6 and Table S2).

**Figure 4.**
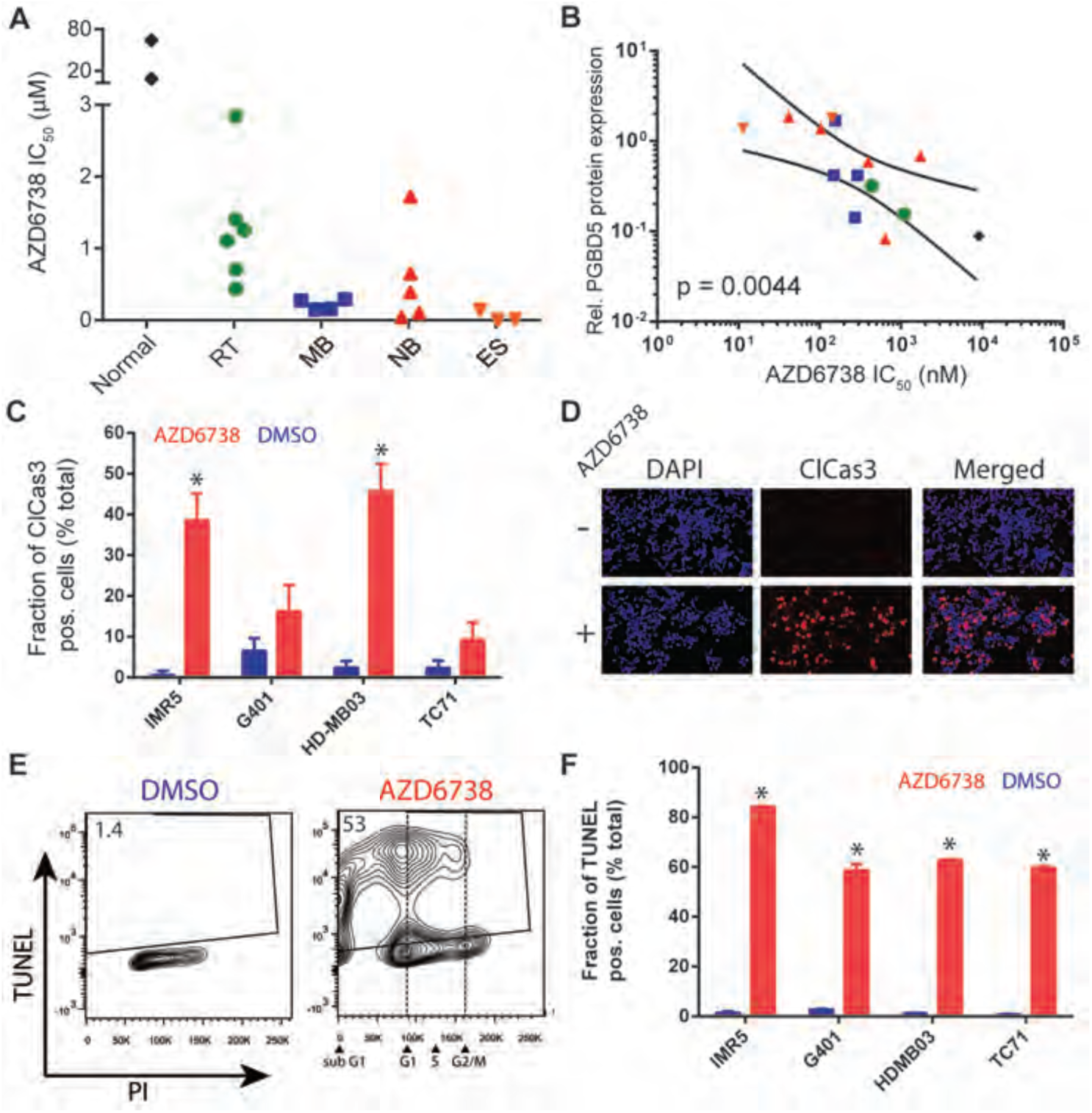
PGBD5-expressing rhabdoid tumor, medulloblastoma, neuroblastoma, and Ewing sarcoma cells exhibit enhanced susceptibility to AZD6738. **(A)** Susceptibility to AZD6738, as shown by its 50% inhibitory concentration (*IC*_*50*_) upon 120 hour treatment of wild-type MEFs and BJ fibroblasts (Normal), rhabdoid tumor (RT), medulloblastoma (MB), neuroblastoma (NB), and Ewing sarcoma (ES) cell lines. Complete list of cell lines and their dose-response growth curves are shown in Figs. S1 & S3. **(B)** PGBD5 protein expression as measured by Western immunoblotting (Fig. S4) is significantly associated with the susceptibility to AZD6738 in pediatric tumor cell lines. Lines denote the 95% confidence interval of linear regression (*p =* 0.0044). Points are labeled according to the color scheme in (A). **(C-F)** Induction of apoptosis and DNA damage, as measured by caspase 3 cleavage (C-D), and TUNEL (E-F) of IMR5 neuroblastoma, G401 rhabdoid tumor, HD-MB03 medulloblastoma, and TC71 Ewing sarcoma cells with 500 nM AZD6738 for 72 hours (red) as compared to DMSO control (blue). \**p* = 6.1 x 10^-4^ and 3.9 x 10^-4^ for AZD6738 versus DMSO, of IMR5 and HD-MB03, respectively. (D) Representative photomicrographs of IMR5 neuroblastoma cells after treatment with 500 nM AZD6738 (+) or DMSO control (-) for 72 hours, stained for cleaved caspase-3 (red) and DAPI (blue). Scale bar = 100 μm. (E) Representative flow cytometric profile of TUNEL and propidium iodide incorporation into HD-MB03 cells after treatment with 500 nM AZD6738 or DMSO control for 48 hours. (F) Induction of TUNEL upon treatment with 500 nM AZD6738 (red) versus DMSO control (blue) for 48 hours. \**p* = 0.042, 0.025, 5.17 x 10^-9^, and 1.98 x 10^-5^ for IMR5, G401, HDMB03 and TC71, respectively. Error bars represent standard deviations of three biologic replicates.

In agreement with observations of MEFs and RPE cells expressing PGBD5, human tumor cell lines expressing endogenous PGBD5 and treated with AZD6738 underwent apoptosis and accumulated unrepaired DNA damage, as measured by caspase 3 cleavage and TUNEL staining (Figs. 4C-F). Most of the TUNEL incorporation induced by AZD6738 in PGBD5-expressing tumor cells was observed in G1 cells (Figs. 4E & S7). Thus, AZD6738 exhibits anti-tumor efficacy against PGBD5-expressing childhood solid tumors *in vitro*.

### Endogenous PGBD5 is necessary to confer enhanced susceptibility of tumor cells to inhibitors of DNA damage signaling

Since ectopic PGBD5 expression may induce DNA damage and signaling dependencies not present in human tumors with endogenous PGBD5, we sought to determine if endogenous PGBD5 expression is required for the enhanced susceptibility to inhibitors of DNA damage signaling. To achieve this, we identified two independent lentiviral RNA interference vectors expressing short hairpin RNA (shRNA) against human *PGBD5* that substantially depleted endogenous PGBD5 both at the mRNA and protein levels, as compared to control vectors targeting *GFP* (Fig. 5A) (*17*). The degree of PGBD5 depletion appeared to depend on tumor type, with rhabdoid G401 cells exhibiting more than 15-fold reduction in mean *PGBD5* levels, whereas neuroblastoma IMR5 and medulloblastoma HD-MB03 cells exhibited reduced effects (Fig. 5A). We found that depletion of *PGBD5* induced significant resistance to AZD6738 as compared to wild-type cells or cells transduced with control vector targeting GFP (shGFP), as evidenced by the relative increase in the *IC*_*50*_ values of rhabdoid tumor G401, neuroblastoma IMR5, and medulloblastoma HD-MB03 cells (Fig. 5B-D, Table S3). Likewise, depletion of endogenous PGBD5 also conferred relative resistance to the ATM-selective inhibitor KU60019, supporting the notion that PGBD5 induces intrinsic DNA damage and requires ongoing DNA damage signaling in tumor cells (Figure S8). Notably, PGBD5-dependent resistance to inhibition of DNA damage signaling appeared to depend on tumor type, consistent with the notion that intrinsic differences in DNA repair and DNA damage signaling may impact therapeutic effects of targeting PGBD5-induced DNA repair dependencies. Thus, PGBD5 expression is necessary to confer susceptibility to inhibitors of DNA damage signaling such as AZD6738 in diverse childhood solid tumors.

**Figure 5.**
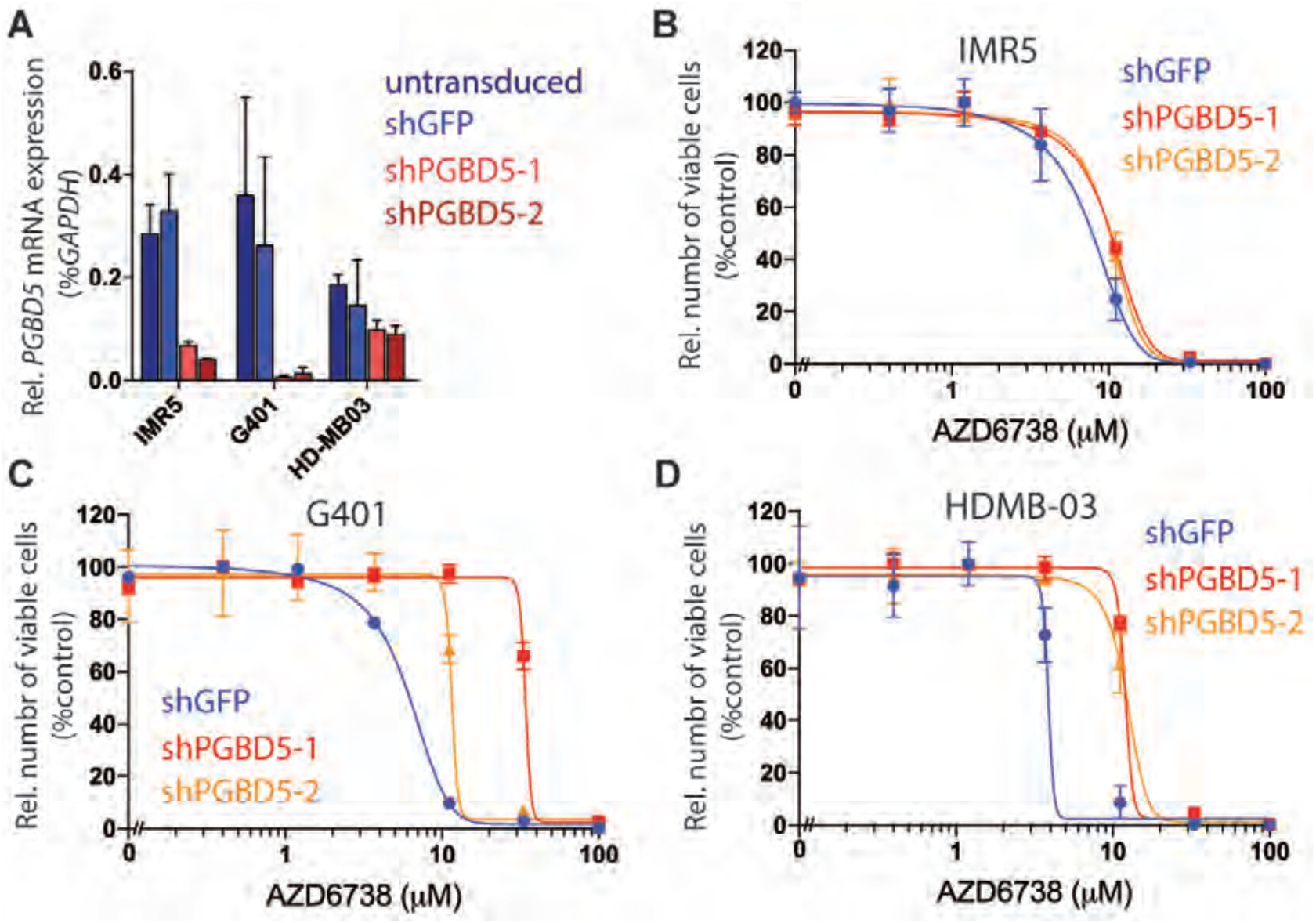
PGBD5 expression is necessary for tumor cell susceptibility to AZD6738. **(A)** *PGBD5* mRNA expression in cells transduced with shRNA against PGBD5, as compared to shGFP and untransduced control cells. **(B-D)** Dose-response of childhood tumor cell lines HDMB-03 (B), IMR5 (C) and G401 (D) treated with varying concentrations of AZD6738 for 120 hours upon *PGBD5* depletion. Error bars represent standard deviations of three replicates.

### AZD6738 induces DNA damage, apoptosis, and exhibits anti-tumor efficacy in xenograft models of high-risk human neuroblastoma and medulloblastoma *in vivo*

Compelled by the finding that AZD6738 induced PGBD5-dependent DNA damage and apoptosis in pediatric solid tumor cell lines *in vitro*, we set out to test whether single-agent AZD6738 treatment has anti-tumor activity in preclinical models of pediatric solid tumors *in vivo*. First, we chose to investigate its activity against mouse xenografts of high-risk *MYCN*-amplified neuroblastoma (IMR5), high-risk MYC group 3 medulloblastoma (HD-MB03), refractory rhabdoid tumor (G401) and Ewing sarcoma (TC-71) cells, as they represent the most common refractory childhood solid tumors (*33, 34*). Thus, we transplanted IMR5, HD-MB03, G401, TC-71 cells subcutaneously in athymic nude *Foxn1*^*nu*^ immunodeficient mice, and monitored tumor growth upon oral treatment of mice with 50 mg/kg/day of AZD6738 (Fig. 6). We found that AZD6738 significantly impaired the growth of both neuroblastoma IMR5 and medulloblastoma HD-MB03 tumors, as compared to vehicle control treated mice *in vivo* (*p* = 4.9e-3 and 5.5e-6 at day 28 respectively, Fig. 6A & B). We did not observe significant toxicity of this treatment, as evidenced by the unchanged animal body weights (Fig. S9). The magnitude of this effect appeared as substantial as compared to the previously reported effects of AZD6738 against tumors with genetic deficiencies of *ATM*, *XRCC1* or *ERCC1* (*35-38*). Similar effects were observed using Kaplan-Meier survival analysis (log-rank *p* = 1.0e-3 and 1.0e-4, respectively, Figure S11). Single-agent anti-tumor activity of AZD6738 against G401 and TC-71 cells were less pronounced *in vivo*, suggesting that tumor-specific differences in DNA repair and DNA damage signaling may impact therapeutic targeting of PGBD5-induced DNA repair dependencies (Fig. S6). Residual tumor cells isolated from mice upon the completion of 20 days of AZD6738 treatment exhibited significantly reduced proliferation, as measured by Ki67 staining (*p* = 3.1e-5 and 1.0e-3 for IMR5 and HD-MB03 respectively, Fig. 6C & D), and increased DNA damage and apoptosis, as measured by γH2AX and cleaved caspase 3 staining, respectively (*p* = 1.4e-3, 4.3e-4, 1.0e-3 and 4.9e-4, respectively, Figs. 6C, E, F). In addition, we also assessed the activity of VE-822, an ATR/ATM inhibitor that is currently in clinical trials (*39*), against neuroblastoma IMR5 and rhabdoid G401 cell line xenografts. In contrast to the ATR-selective AZD6738, single-agent treatment with VE-822 had no significant effects on tumor growth *in vivo* (Fig. S11), supporting the notion that pharmacologic and selectivity properties of inhibitors of DNA damage signaling may also affect the efficacy of PGBD5-dependent therapeutic targeting.

**Figure 6.**
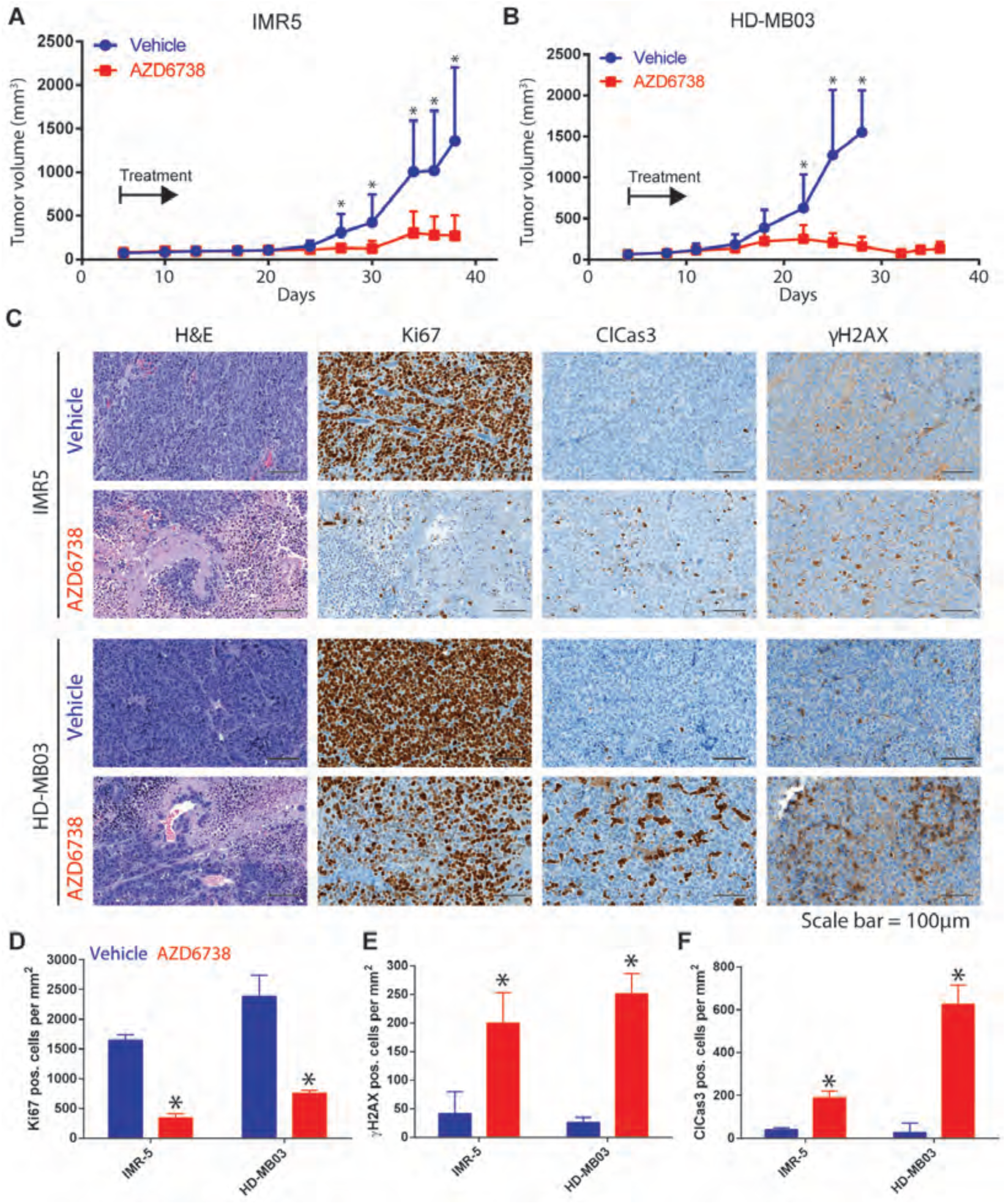
AZD6738 induces DNA damage, apoptosis, and exhibits anti-tumor efficacy in xenograft models of high-risk medulloblastoma and neuroblastoma *in vivo.* **(A-B)** Tumor volumes over time of nude mice harboring IMR5 (A) and HD-MB03 (B) subcutaneously xenografted tumors treated with AZD6738 by oral gavage at 50 mg/kg body weight/day (red) or as compared to vehicle control (blue). Asterisks denote *p* < 0.05. Error bars represent standard deviations of 10 individual xenograft mice per group. Arrows denote start of treatment. **(C)** Representative photomicrographs of sections from IMR5 (top) or HD-MB03 (bottom) tumors upon completion of treatment with AZD6738 (50 mg/kg/day) or vehicle control *in vivo*, and stained for hematoxylin and eosin H&E, Ki67, cleaved caspase-3, and γH2AX, as indicated. Scale bar = 100 μm. **(D-F)** Quantification of the number of cells positively stained for Ki67 (D), cleaved caspase-3 (E) and γH2AX (F) in IMR5 or HD-MB03 xenograft tumors upon completion of treatment with AZD6738 (50 mg/kg/day, red) or vehicle control (blue) *in vivo*. \**p* = 3.1 x 10^-5^ and 0.001 for Ki67 in AZD6738 versus vehicle-treated IMR5 and HD-MB03, respectively. \**p* = 0.001 and 4.9 x 10^-4^ for cleaved caspase-3 in AZD6738 versus vehicle-treated IMR5 and HD-MB03, respectively. \**p* = 0.014 and 4.3 x 10^-4^ for γH2AX in AZD6738 versus vehicle-treated IMR5 and HD-MB03, respectively. Error bars represent standard deviations of 3 independent fields analyzed.

### Synergistic targeting of PGBD5-induced DNA repair dependency in patient-derived primary neuroblastoma xenografts *in vivo*

Considering that AZD6738 exhibited potent single-agent activity against high-risk neuroblastoma and medulloblastoma cell line xenografts, with reduced apparent activity against rhabdoid and Ewing sarcoma xenografts, we reasoned that specific agents that induce DNA damage may be used to selectively potentiate the anti-tumor effects of AZD6738. To test this hypothesis, we analyzed the combination of AZD6738 with cisplatin, a chemotherapeutic drug that crosslinks DNA and is frequently used to treat childhood solid tumors (*40, 41*) (Fig. 7). We found significant synergy between cisplatin and AZD6738 at all drug concentrations tested, as indicated by their low combination indices for the neuroblastoma, medulloblastoma, rhabdoid tumor, and Ewing sarcoma cell lines (*42*) (Fig. 7A).

**Figure 7.**
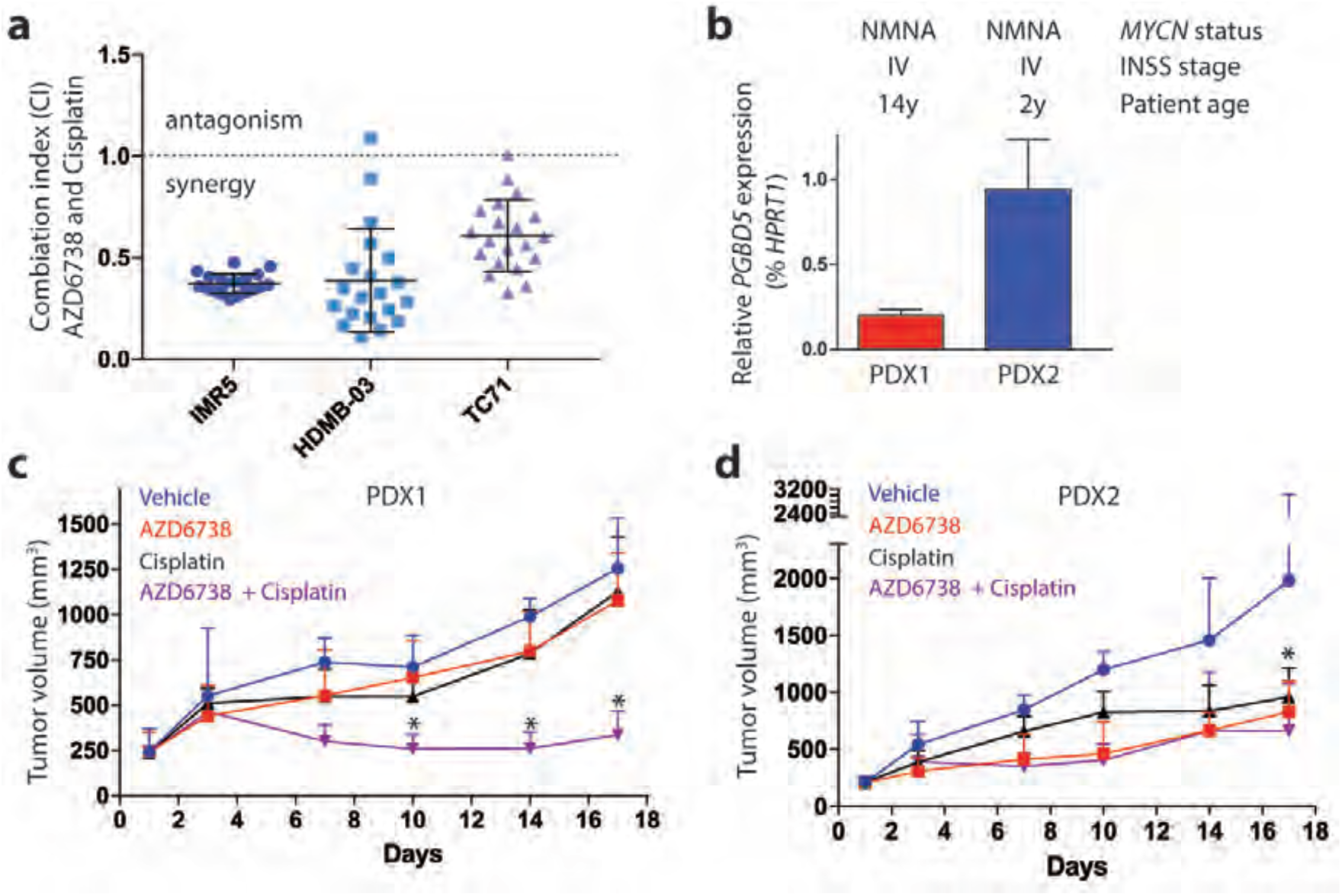
Synergistic targeting of PGBD5-induced DNA repair dependency in primary patient derived high-risk neuroblastoma xenografts in vivo. **(A)** Combination indices (CI) for IMR5, HDMB-03 and TC71 cells treated with cisplatin and AZD6738. **(B)** *PGBD5* mRNA expression in two patient derived high-risk neuroblastoma xenografts; complete demographic and molecular features are described in Table S4 **(C)** Tumor volumes over time of mice engrafted with PDX1 and treated with 50 mg/kg AZD6738 by daily oral gavage (red), as compared to vehicle control (blue), 2 mg/kg cisplatin by weekly IP (black), or combination of AZD6738 and cisplatin (violet). **(D)**Tumor volumes over time of mice engrafted with PDX2 and treated with 50 mg/kg AZD6738 by daily oral gavage (red), as compared to vehicle control (blue), 2 mg/kg cisplatin by weekly IP (black), or combination of AZD6738 and cisplatin (violet). Asterisks denote *p* < 0.05. Error bars represent standard deviations of 4 individual xenograft mice per group.

To investigate the potential therapeutic benefit of combining AZD6738 with cisplatin therapy as a prelude to its clinical testing in patients, we established two patient-derived primary xenografts PDX1 and PDX2 from non-*MYCN* amplified, stage IV metastatic neuroblastoma tumors obtained at diagnosis prior to therapy (Figure 7B, Table S4). In agreement with results in human tumor cell lines (Fig. 4), we observed varying levels of *PGBD5* expression in the two neuroblastoma xenografts (Fig. 7B). Subsequently, we transplanted these tumor specimens subcutaneously into NOD.Cg-*Prkdc*^*scid*^ *Il2rg*^*tm1Sug*^/JicTac (NOG) immunodeficient mice, and upon tumor engraftment as evidenced by tumor volumetric measurements, randomized recipient mice to be treated with single-agent AZD6738 (50 mg/kg daily PO for 14 days), single-agent cisplatin (2 mg/kg IP every 7 days), combination of AZD6738 and cisplatin, or vehicle control (Fig. 7C & D). Consistent with the relatively low level of *PGBD5* expression in PDX1 (Fig. 7B), we found that single-agent treatment with AZD6738 or cisplatin had limited effects, whereas combination of AZD6738 with cisplatin exhibited significant reduction in tumor growth, as compared to single-agent or vehicle control treatments (*p* = 1.9e-3, Fig. 7C). Likewise, for PDX2 that had relatively high *PGBD5* expression (Fig. 7B), we observed significant reduction in tumor growth upon single-agent AZD6738 and cisplatin treatment, as compared to vehicle control (*p* = 0.032 and 0.079, respectively, Fig. 7D). Altogether, these results indicate that AZD6738 exhibits significant single-agent and cisplatin-combination efficacy against PGBD5-expressing solid tumors.

## Discussion

We have now found that the PGBD5 DNA transposase expressed in the majority of childhood solid tumors confers a synthetic dependency on DNA damage repair and signaling. Consistent with the genomic rearrangements promoted by PGBD5 in rhabdoid tumors (*17*), expression of PGBD5 induces DNA damage, which requires both DNA damage repair and DNA damage signaling, resulting in apoptosis if impaired by their selective inhibitors (Fig. 8). Indeed, both primary mouse and human cells engineered to express PGBD5, as well as PGBD5-expressing childhood solid tumors, accumulated unrepaired DNA damage and underwent apoptosis upon treatment with selective inhibitors of DNA damage signaling. This effect was due to the specific nuclease activity of PGBD5, insofar as mutation of its putative catalytic nuclease residues completely abrogated the dependence on DNA damage signaling. In turn, single-agent treatment with the DNA damage signaling inhibitor AZD6738 exhibited potent anti-tumor activity against high-risk neuroblastomas and medulloblastomas that express high levels of PGBD5 in preclinical mouse models *in vivo*.

**Figure 8.**
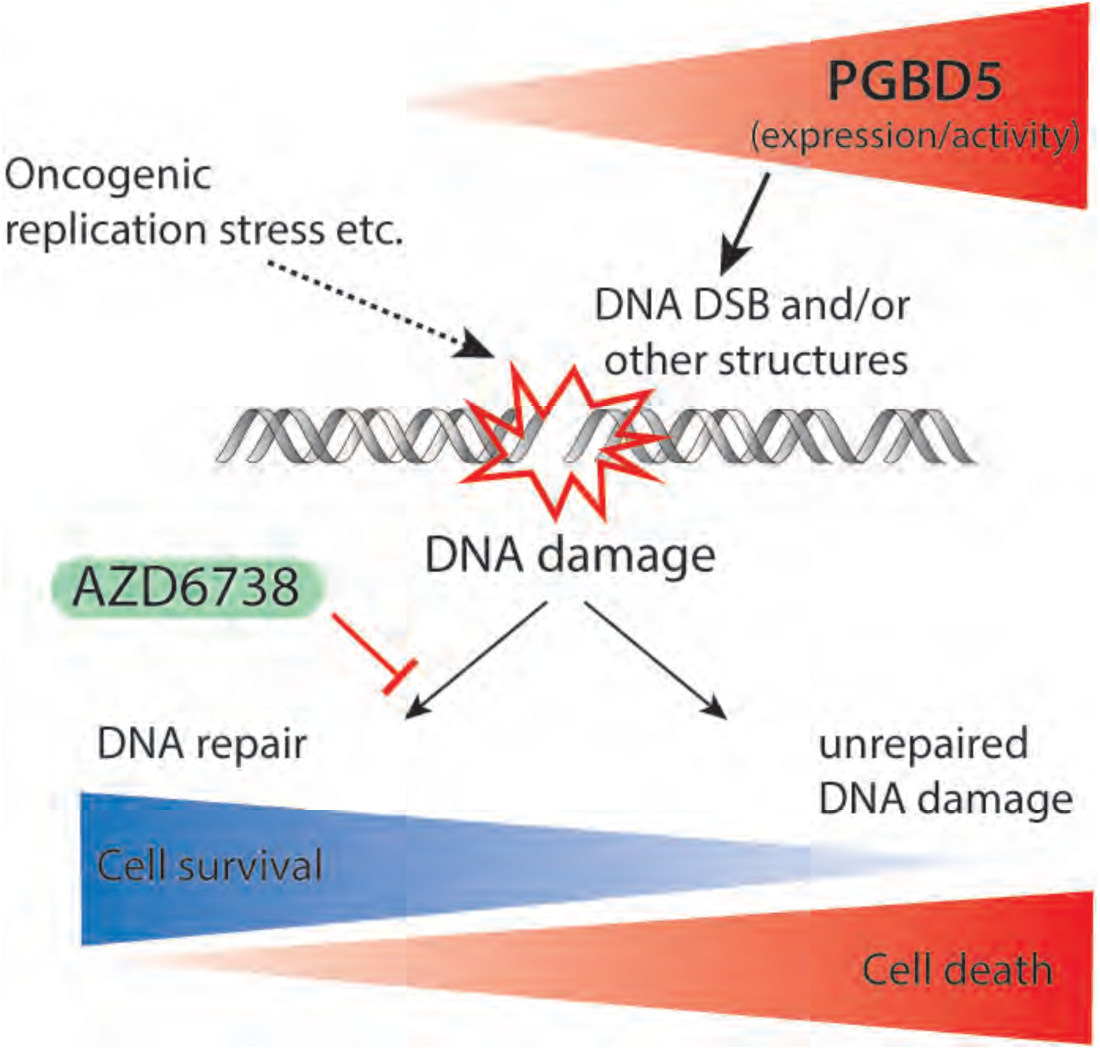
Model of therapeutic targeting of PGBD5-induced DNA damage signaling synthetic dependency. Tumors with increasing levels of PGBD5 expression and DNA recombinase activity accumulate DNA damage, in concert with other intrinsic sources of cellular DNA damage such as replication stress. PGBD5-dependent DNA damage leads to the generation of various DNA damage structure and DNA damage signals, which in turn activate distinct arms of the DNA damage signaling. Consequently, PGBD5-dependent DNA damage signaling can be inhibited using selective pharmacologic inhibitors, inducing accumulation of DNA damage, impairing DNA repair, and leading to cell death.

Human cancers harbor numerous mechanisms of endogenous DNA damage as a source of genetic mutations and requirements for active DNA damage repair (*43*). As a result, selective inhibitors of DNA damage signaling exhibit anti-tumor activities in various cancers (*44-47*). In particular, selective inhibitors of ATR and ATM kinases have been used to target tumors with intrinsic deficiencies in DNA repair (*43*), as well as DNA damage susceptibility, such as that induced by oncogene and replication stress (*47*). Our current work now revealed a specific synthetic dependency conferred by the endogenous DNA transposase PGBD5 in the majority of childhood solid tumors. Genetic experiments showed equivalent functional requirements for the scaffolding KU complex that directly binds DSBs, and the ATR and ATM kinases that mediate DNA damage signaling. However, chemical DNA damage signaling kinase inhibitors exhibited a specific profile with the ATR-and ATM-selective AZD6738 and KU60019 inhibitors exhibiting enhanced PGBD5-dependent activity. Given their varied potency and selectivity, it is possible that other selective DNA damage signaling inhibitors can also be used to effectively target PGBD5-induced tumor synthetic dependencies. Since ATR and ATM are activated by specific DNA structures, including single-stranded DNA bound by RPA (*48*), modified bases (*49*), and chromatin assemblies (*50*), the apparent activity of selective inhibitors of DNA damage signaling may be due to the PGBD5-induced generation of as of yet unrecognized DNA or chromatin structures that recruit specific components of the DNA damage response (*51*). While PGBD5-dependent DNA damage and apoptosis induced by AZD6738 appeared to occur in G1 phase without measureable replication stress or ATM depletion, it is also possible that inhibition of ATR or additional kinases by AZD6738 in S phase leads to apoptosis in the subsequent G1 phase of the cell cycle. Lastly, the susceptibility of PGBD5-expressing tumors to selective inhibitors of DNA damage signaling is expected to cooperate with other sources of DNA damage in tumors, such as oncogene or replication stress.

We anticipate that susceptibility to AZD6738 and other therapeutic strategies to target PGBD5-dependent DNA repair requirements may depend on the tumor-specific mechanisms of DNA repair, including variability in the expression and activity of non-homologous end-joining and homologous recombination, as determined by variation in tumorigenic mechanisms and cells of origin (*15*). For example, neuroblastoma cells can have varying activity of NHEJ repair as partly affected by neurotrophin receptor signaling (*52, 53*), which may impact the therapeutic efficacy of targeting PGBD5-induced DNA repair dependencies. Such a model, in which intrinsic differences in DNA repair and DNA damage signaling impact therapeutic targeting of PGBD5-induced DNA repair dependencies, is supported both by the apparent variation in tumor response to AZD6738 as well as its synergistic potentiation by cisplatin therapy (Fig. 4 & 8).

Finally, our findings may explain the apparent activity of other kinase inhibitors, such as for example the observed anti-tumor activity of dactolisib (NVP-BEZ235) in neuroblastoma, medulloblastoma and Ewing sarcoma (*54-56*), given its potent kinase inhibition of ATR (*12*). Inhibitors of DNA damage signaling are currently being investigated in clinical trials, including AZD6738 (NCT02264678, NCT02223923, NCT02630199). Our findings warrant their immediate investigation in clinical trials for children with solid tumors, the majority of which express PGBD5 and should be susceptible to targeted therapy of this synthetic DNA damage signaling dependency. We anticipate that improved understanding of the molecular synthetic dependencies and their targeting by emerging selective pharmacologic inhibitors should lead to rational therapeutic strategies for refractory solid tumors.

## Material and Methods

### Reagents

All reagents were obtained from Sigma Aldrich if not otherwise specified. Synthetic oligonucleotides were obtained from Eurofins (Eurofins MWG Operon, Huntsville, AL, USA) and purified by HPLC. All small molecule inhibitors used were dissolved in dimethyl sulfoxide at 10 mM (Fisher Scientific, Watham, MA, USA) and stored at −20 °C until further use: AZD6738 (Astra Zeneca, London, UK), KU60019 (Selleck Chemicals LLC, Munich, Germany), AZ20 (Tocris, Bristol, UK), VE822 (Selleck Chemicals LLC), AZD7762 (Selleck Chemicals LLC).

### Plasmid constructs

Human *PGBD5* cDNA (Refseq ID: NM_001258311.1) was cloned into the lentiviral vector in frame with N-terminal GFP to generate pRecLV103-GFP-PGBD5 (GeneCopoeia, Rockville, MD, USA). pReceiver-Lv103 encoding *GFP* was used as a negative control in all experiments. Missense *GFP-PGBD5* mutants were generated using site-directed mutagenesis according to the manufacturer’s instructions (QuikChange Lightning, Agilent, Santa Clara, CA, USA) as described (*16*). Doxycycline-inducible pINDUCER21 vector was obtained from Thomas Westbrook (*57*), and used to generate pINDUCER21-PGBD5 using Gateway cloning, according to the manufacturer’s instructions (Fisher Scientific, Watham, MA, USA). pLKO.1 shRNA vectors targeting PGBD5 (TRCN0000138412, TRCN0000135121) and control shGFP were obtained from the RNAi Consortium (Broad Institute, Cambridge, MA).

### Lentivirus production and cell transduction

Lentivirus production was carried out as previously described (*58*). Briefly, HEK293T cells were transfected using TransIT-LT1 with 2:1:1 ratio of the lentiviral vector, and psPAX2 and pMD2.G packaging plasmids, according to manufacturer’s instructions (Mirus, Madison, WI, USA). Virus supernatant was collected at 48 and 72 hours post-transfection, pooled, filtered and stored at −80 °C. RPE and BJ cells were transduced with pRecLV103 virus particles at a multiplicity of infection (MOI) of 5 in the presence of 8 μg/ml hexadimethrine bromide. Transduced cells were selected for 2 days with puromycin hydrochloride (RPE cells at 10 μg/ml, BJ cells at 2 μg/ml) or G418 sulfate (2 mg/ml), depending on the vector-mediated resistance. For pINDUCER21 viruses, cells were transduced at a MOI of 1, and isolated using fluorescence-activated cell sorting (FACSAria III, BD Bioscience, San Jose, CA, USA). For lentiviral transduction of shRNAs, cells were selected for 2 days with puromycin hydrochloride at 2 μg/ml.

### Cell culture

All cell lines were obtained from the American Type Culture Collection if not otherwise specified (ATCC, Manassas, Virginia, USA). *Ku80*^*-/-*^, *Atr*^*S/S*^, and *Atm*^*-/-*^ mouse embryonic fibroblasts were generated by transduction with SV40 large T antigen (*24-26*). All neuroblastoma cell lines were provided by Johannes H. Schulte. HD-MB03 cells were generated by Milde et al. (*59*). Rhabdoid tumor cell lines KP-MRT-NS, KP-MRT-YM, KP-MRT-RY and MP-MRT-AN were generated by Yasumichi Kuwahara and Hajime Hosoi. The identity of all cell lines was verified by STR analysis (Genetica DNA Laboratories, Burlington, NC, USA) and absence of *Mycoplasma sp.* contamination was determined using Lonza MycoAlert (Lonza Walkersville, Inc., Walkersville, MD, USA). Cell lines were cultured in 5% CO_2_ in a humidified atmosphere at 37 °C in Dulbecco’s Modified Eagle medium with high glucose (DMEM-HG) supplemented with 10 % fetal bovine serum (FBS) and antibiotics (100 U / ml penicillin and 100 μg / ml streptomycin). Cell viability was assessed using CellTiter-Glo according to the manufacture’s protocol (Promega, Madison, WI, USA).

### Quantitative RT-PCR

RNA was isolated using RNeasy Mini, according to manufacturer’s instructions (Qiagen, Venlo, Netherlands). cDNA was synthesized using the SuperScript III First-Strand Synthesis System, according to the manufacturer’s instructions (Invitrogen, Waltham, MA, USA). Quantitative real-time PCR was performed using the KAPA SYBR FAST PCR polymerase with 20 ng template and 200 nM primers, according to the manufacturer’s instructions (Kapa Biosystems, Wilmington, MA, USA). PCR primers are listed in Table S5. Ct values were calculated using ROX normalization using the ViiA 7 software (Applied Biosystems).

### Western blotting

To analyze protein expression by Western immunoblotting, 1 million cells were suspended in 80 μl of lysis buffer (4% sodium dodecyl sulfate, 7% glycerol, 1.25% beta-mercaptoethanol, 0.2 mg/ml Bromophenol Blue, 30 mM Tris-HCl, pH 6.8) and incubated at 95 °C for 10 minutes. Cell suspensions were lysed using Covaris S220 adaptive focused sonicator, according to the manufacturer’s instructions (Covaris, Woburn, CA). Lysates were cleared by centrifugation at 16,000 g for 10 minutes at 4 °C. Clarified lysates (30 μl) were resolved using sodium dodecyl sulfate-polyacrylamide gel electrophoresis, and electroeluted using the Immobilon FL PVDF membranes (Millipore, Billerica, MA, USA). Membranes were blocked using the Odyssey Blocking buffer (Li-Cor, Lincoln, Nebraska, USA), and blotted using antibodies listed in Table S6. Blotted membranes were visualized using goat secondary antibodies (Table S6) conjugated to IRDye 800CW or IRDye 680RD and the Odyssey CLx fluorescence scanner, according to manufacturer’s instructions (Li-Cor, Lincoln, Nebraska, USA). Image analysis was done using the Li-Cor Image Studio software (version 4). Relative protein expression was calculated as fluorescence intensity of the protein of interest relative to the fluorescence intensity of the loading control protein. At least 3 blots were analyzed for each condition.

### Flow cytometry

Cells were fixed using neutral-buffered formalin for 10 min on ice, washed with PBS, resuspended in 0.1% Triton-X100 in PBS, and incubated for 15 min at room temperature. Permeabilized cells were washed twice with PBS, and resuspended in 100 μl of Hank’s balanced salt solution (HBSS) with 0.1% bovine serum albumin and 2 μl of Alexa Fluor 647-conjugated antibody against cleaved caspase-3 (Table S6). Cells were incubated for 30 min at room temperature in the dark washed twice with PBS and stained with 1 μg/ml DAPI. Cells were analyzed on the Fortessa LSR as described before (BD Bioscience) (*60, 61*). TUNEL staining was done using the APO-BrdU TUNEL Assay Kit, according to the manufacturer’s protocol (Fisher Scientific). EdU labeling was done using the EdU Click-iT kit according to the manufacture’s protocol (Fisher Sceintific).

### Histological staining

Histologic processing and staining was done as described previously (*62, 63*). In short, cells were plated on 8-well glass Millicell EZ chamber slides at 5000 cells/well, grown for 24 hours, and fixed using 4% paraformaldehyde for 10 min at room temperature (Millipore). Tumor xenograft tissue was fixed using 4% paraformaldehyde for 24 hours at room temperature. Tissues were embedded in paraffin using the ASP6025 tissue processor (Leica, Wetzlar, Germany), sectioned at 5 μm using the RM2265 microtome (Leica), and collected on SuperfrostPlus slides (Fisher Scientific). Tissue sections were deparaffinized with EZPrep buffer (Ventana Medical Systems). Antigen retrieval was performed with Cell Conditioning 1 buffer (Ventana Medical Systems), and sections were blocked for 30 minutes with Background Buster solution (Innovex, Norwood, MA, USA). Primary antibodies were applied for 5 hours at 1 μg/ml (Table S6). Secondary antibodies were applied for 60 minutes (Table S6).

For immunohistochemistry staining, diaminobenzidine (DAB) detection was performed with the DAB detection kit according to manufacturer instruction (Ventana Medical Systems). Slides were counterstained with hematoxylin and a cover slip was mounted with Permount (Fisher Scientific).

For immunofluorescence staining, the detection was performed with Streptavidin-HRP D (Ventana Medical Systems), followed by incubation with Tyramide Alexa Fluor 647 prepared according to manufacturer instruction (Invitrogen). Slides were then counterstained with 5 μg/ml DAPI for 10 min and a cover slip was mounted with Mowiol (Sigma Aldrich). TUNEL and Hoechst staining was done using Click-iT TUNEL Alexa Fluor Imaging Assay according to the manufacture’s protocol.

### Image acquisition and analysis

Bright-field images were acquired on an Axio Observer microscope (Carl Zeiss Microimaging, Oberkochen, Germany). Epifluorescence images were acquired using the EVOS FL microscope (Thermo Fisher). Slides were scanned using the Pannoramic 250 slide scanner and images analyzed using the Pannoramic Viewer (3DHistech, Budapest, Hungary). At least 100 stained cells were counted in at least 3 independent fields of view.

### Xenografts

All mouse experiments were carried out in accordance with institutional animal protocols. Five million cells were suspended in 200 μl Matrigel (BD Bioscience, Heidelberg, Germany) and injected subcutaneously into the left flank of 6-week-old female athymic nude *Foxn1*^*nu*^ mice (The Jackson Laboratory, Bar Harbor, Maine, USA). Patient-derived xenografts were performed using NOD.Cg-*Prkdc*^*scid*^ *Il2rg*^*tm1Sug*^/JicTac (NOG) mice (Taconic). Patient tumors were serially transplanted in mice three times before experiments. Tumor growth was monitored using caliper measurements, and tumor volume was calculated using the formula 3.14159 x length x width ^2^ / 6000. Mice were sacrificed by CO_2_ asphyxiation when tumor size exceeded 2,000 mm^3^. Mice were treated with AZD6738 at 50 mg/kg body weight once a day per oral gavage. For *in vivo* treatment, AZD6738 was dissolved in DMSO at 62.5 mg/ml and mixed 1:10 in 40% propylene glycol and 50% sterile water resulting in a final AZD6738 concentration of 6.25 mg/ml. VE-822 was dissolved in 10% Vitamin E d-alpha tocopheryl polyethylene glycol 1000 succinate and administered by oral gavage. Cisplatin was dissolved in phosphate buffered saline and administered intraperitoneally at 2 mg/kg body weight once every 7 days.

### Statistical analysis

For comparisons between two sample sets, statistical analysis of means was performed using 2-tailed, unpaired Student’s t-tests. Survival analysis was done using the Kaplan-Meier method, as assessed using a log-rank test.

## Acknowledgments

We thank Alejandro Gutierrez, Marc Mansour, Michael Kharas, and John Poirier for critical discussions, and Wong Wai for technical assistance. We thank the Experimental Pharmacology & Oncology Berlin-Buch GmbH for assistance with patient-derived xenografts.

## Author Contributions

AGH: study design, collection and interpretation of the data; CR, HDG, IM, JVS, PH, EJ, JK, WW, EDS, YK, HH, NG, AK, JP, JS: data collection; AK: study design and data analysis. AGH and AK wrote the manuscript with contributions from all authors.

## Data and materials availability

Data are openly available at the Zenodo digital repository (https://zenodo.org/).

## Supplementary Materials

**Henssen et al. Therapeutic targeting of PGBD5-induced DNA repair dependency in pediatric solid tumors.**

## Supplementary Figures

**Figure S1.**
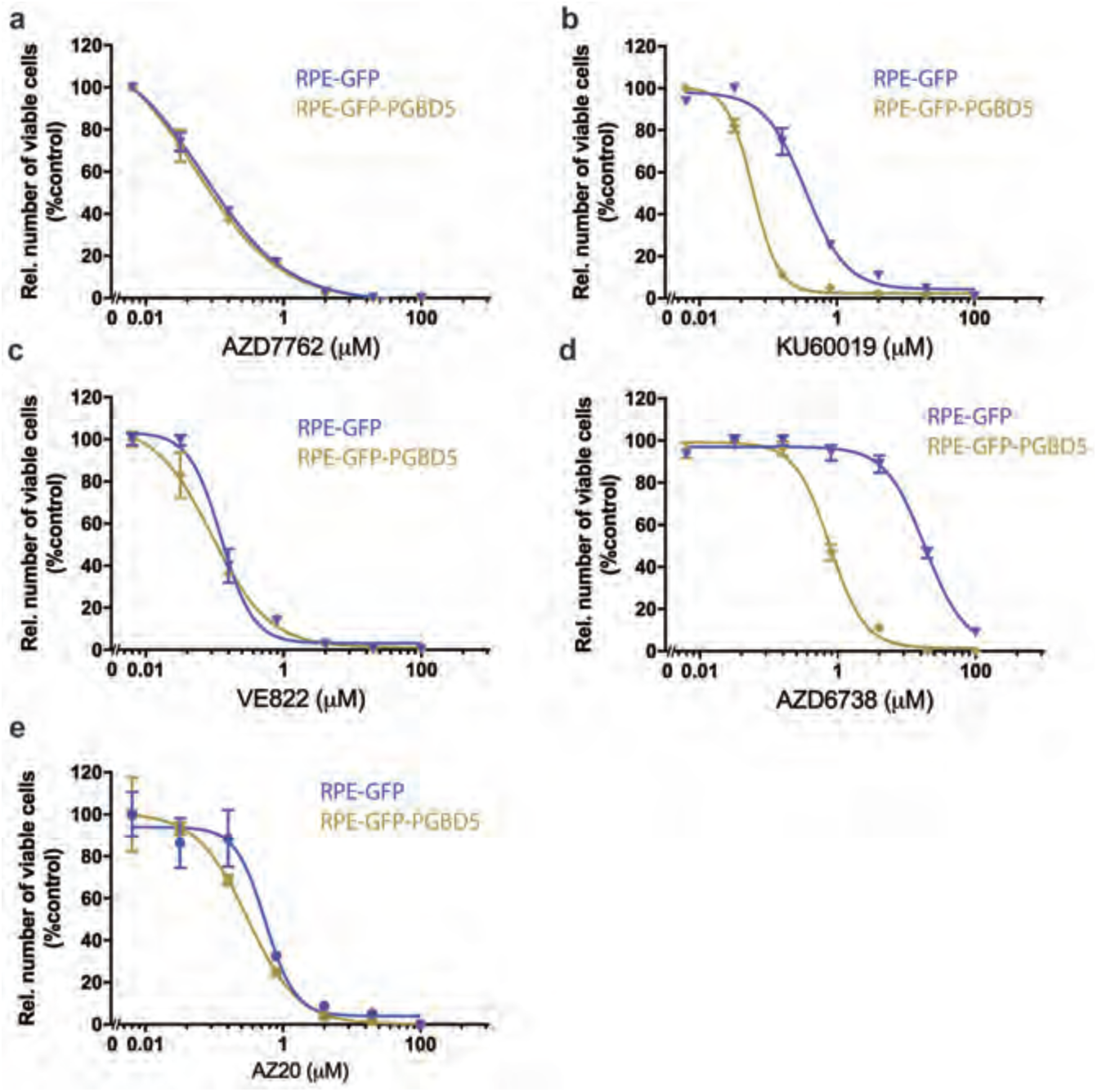
PGBD5 expression is sufficient to confer sensitivity to ATM and ATR inhibitors. Dose response curves of RPE cells expressing *GFP-PGBD5* or *GFP* control treated with AZD7762 (A), KU60019 (B), VE822 (C), AZD6738 (D) and AZ20 (E) for 120 hours. Error bars indicate standard deviation of three biologic replicates.

**Figure S2.**
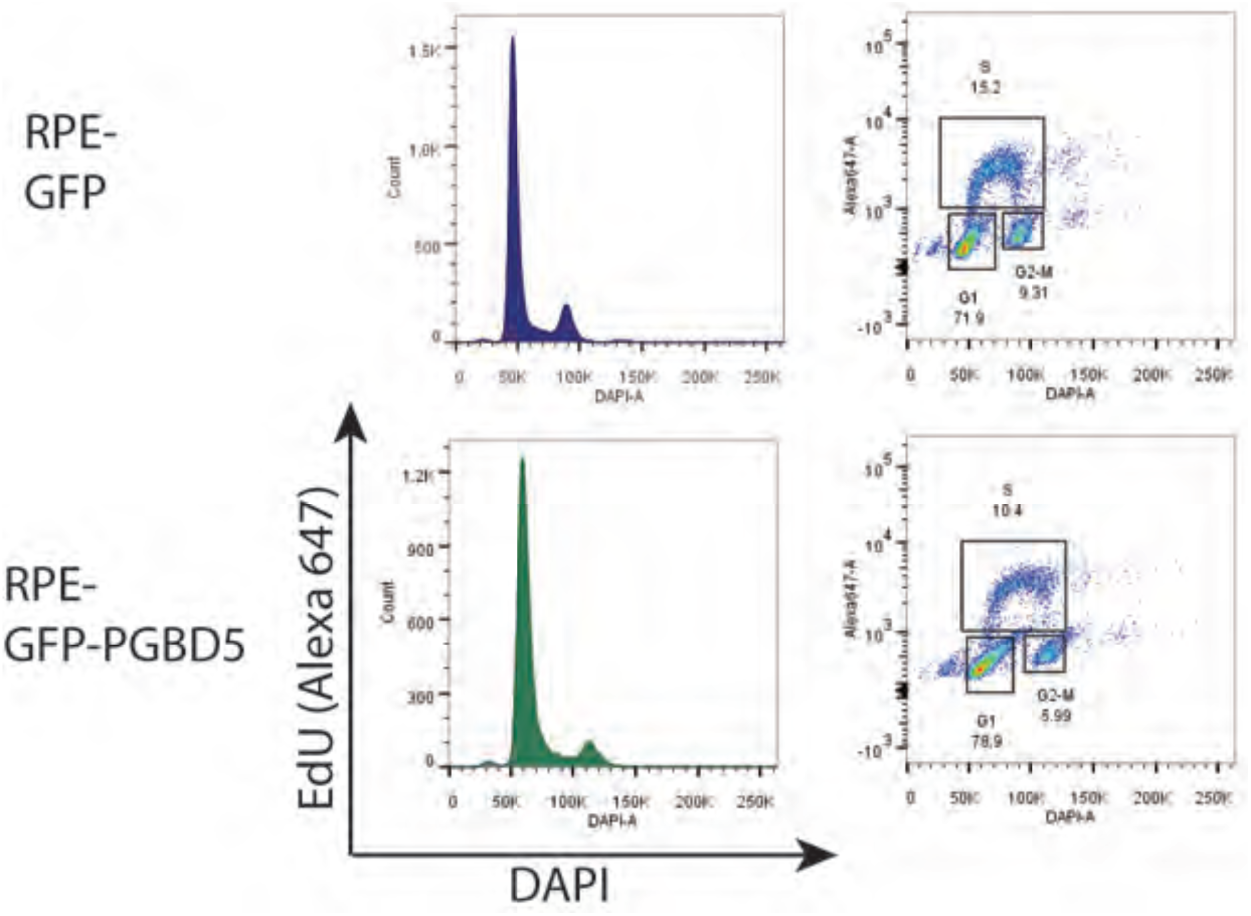
GFP-PGBD5 expression in RPE cells does not increase cell cycling. FACS plot of RPE cells expressing GFP-PGBD5 as compared to cells expressing GFP control, labeled with EdU Alexa 647 and DAPI.

**Figure S3.**
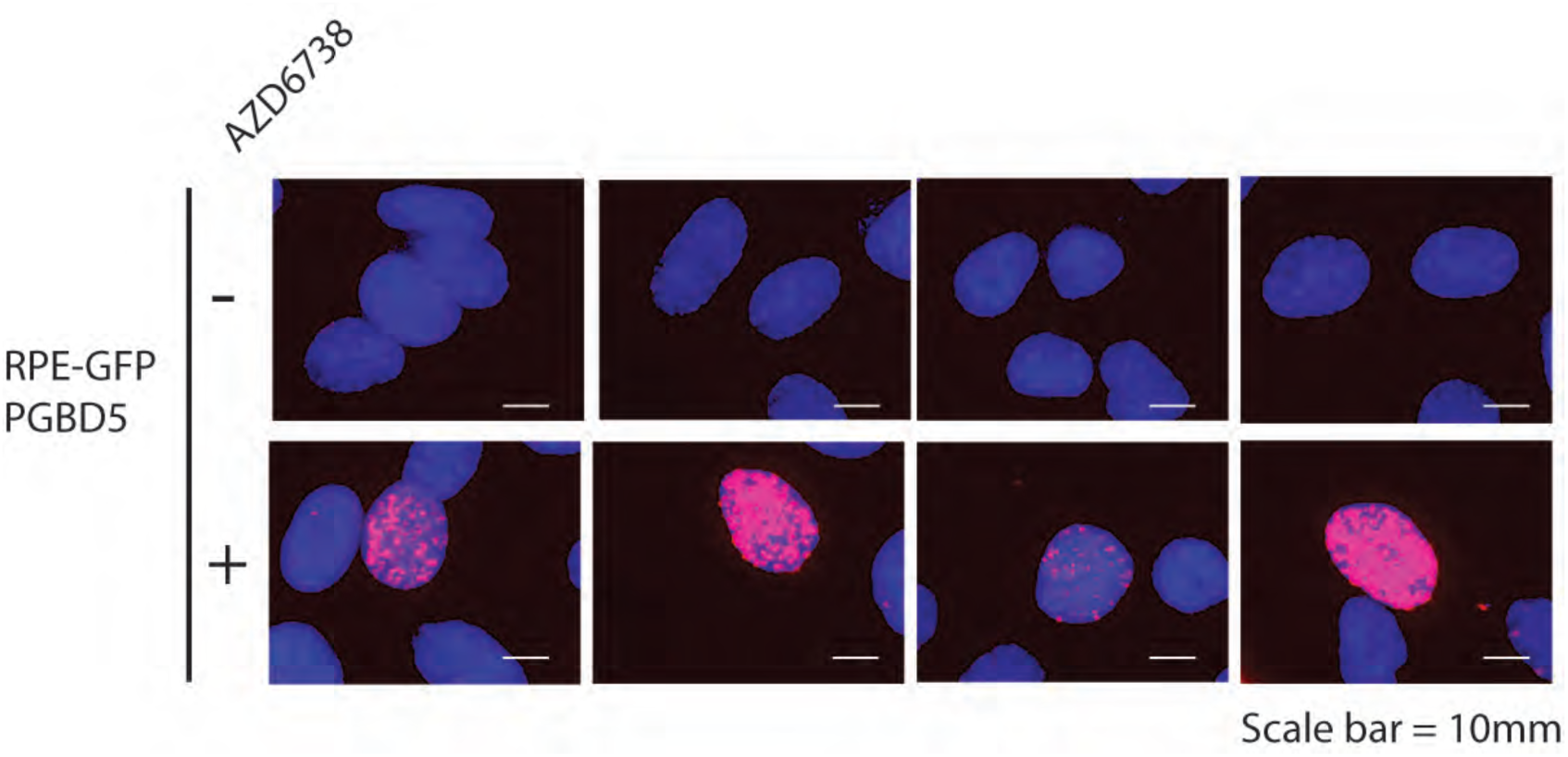
AZD6732 induces punctate γH2AX accumulation. Representative photomicrographs of RPE cells expressing GFP-PGBD5 and treated with AZD6732.

**Figure S4.**
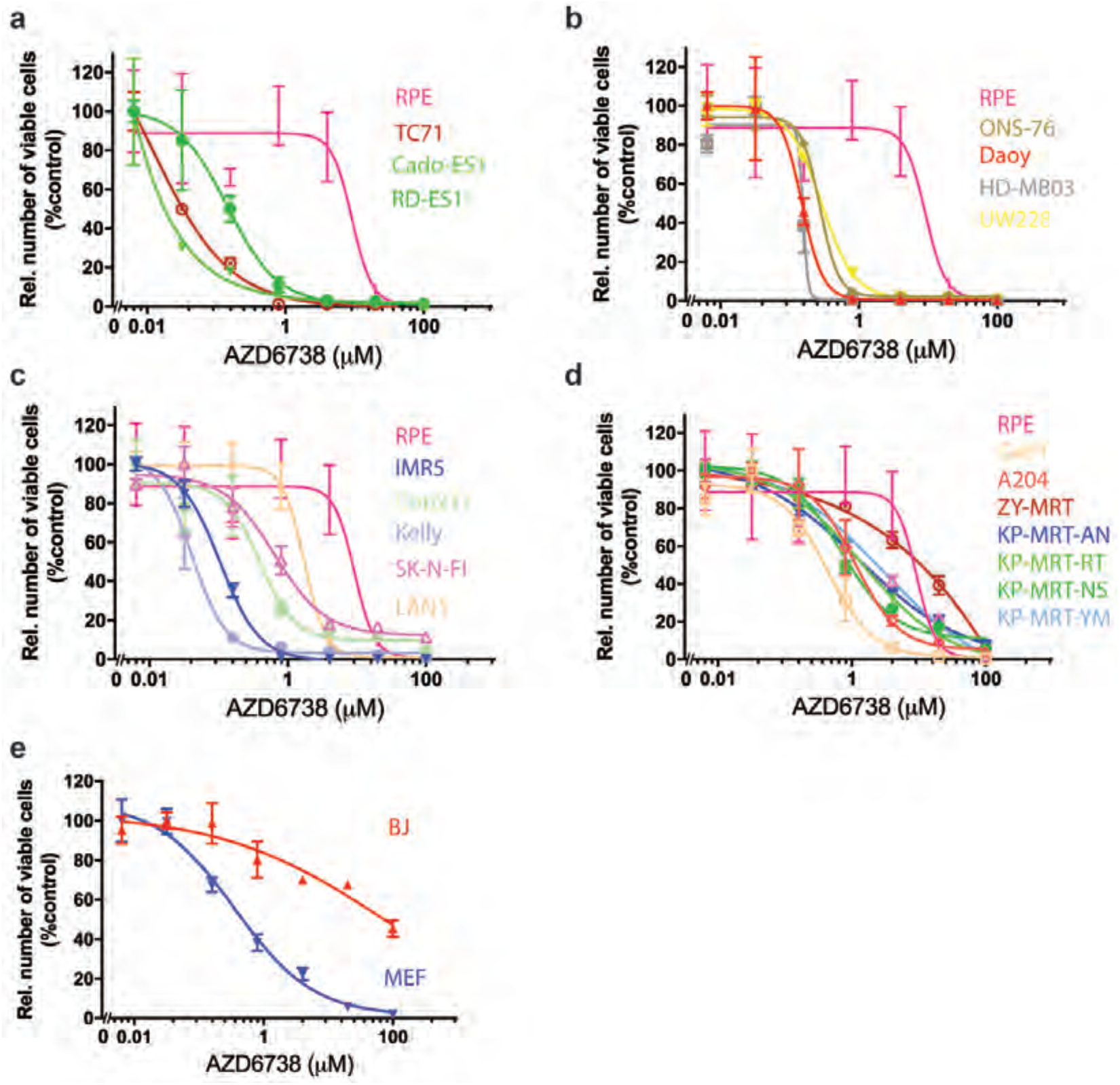
Rhabdoid tumor, medulloblastoma, neuroblastoma, and Ewing sarcoma cells exhibit sensitivity to AZD6738. Dose response curves of Ewing sarcoma (A), medulloblastoma (B), neuroblastoma (C), rhabdoid tumor cells (D) and non-transformed human cells (E) treated with AZD6738 for 120 hours. Error bars indicate standard deviation of three biologic replicates.

**Figure S5.**
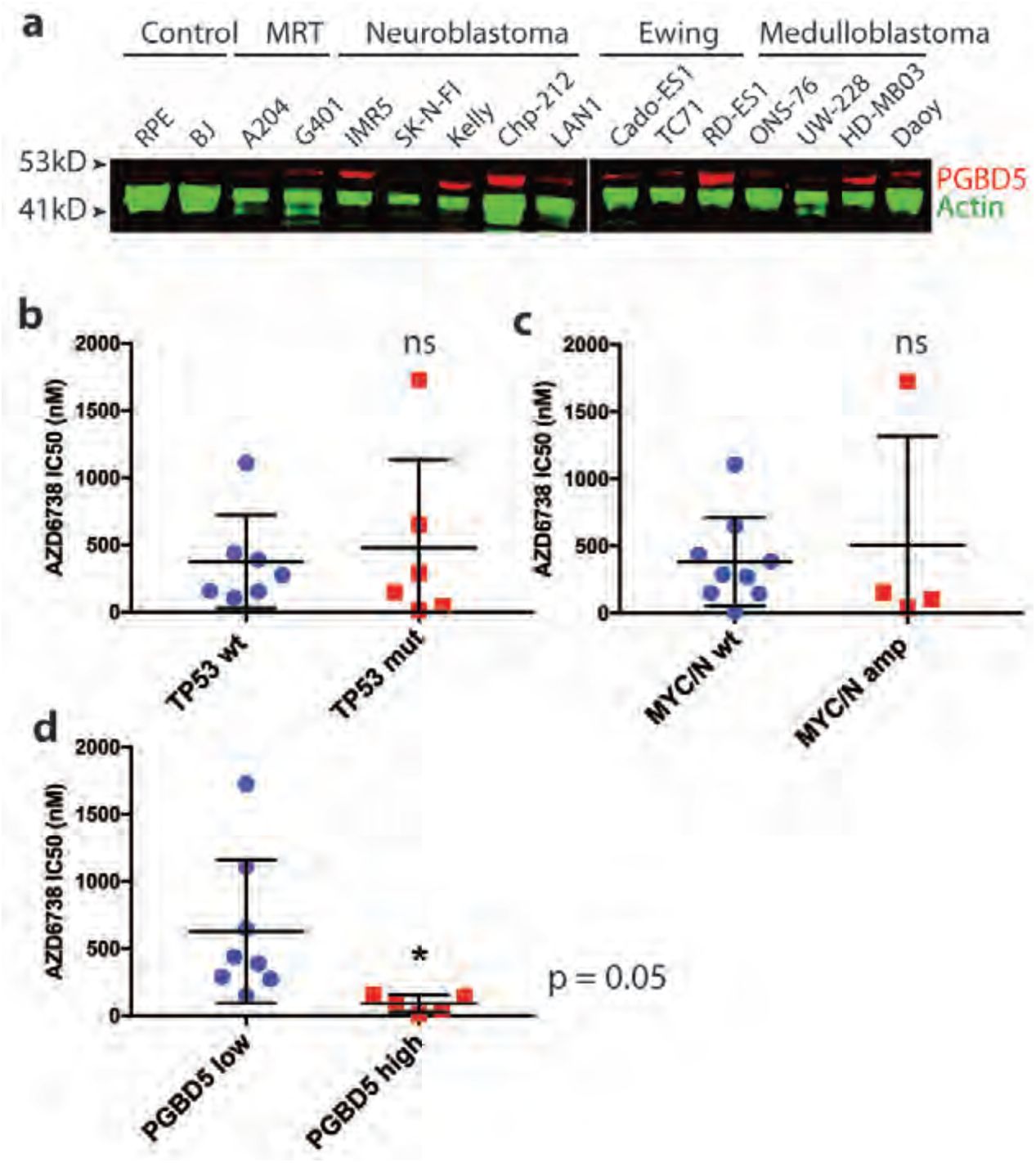
Expression of PGBD5 in tumor cell lines significantly correlates to their susceptibility to AZD6738 whereas *TP53* and *MYC/N* status do not. **(A)** Quantitative fluorescence Western immunoblotting of PGBD5 in tumor and normal cells. Actin serves as the loading control. **(B)** Inhibitory concentration 50% values (IC_50_) for AZD6738 in *TP53* wild-type as compared to *TP53-*mutant pediatric tumor cell lines (unpaired t-test, *p* > 0.05 for *TP53* mut. compared to *TP53* wt., not significant, ns). **(C)** IC_50_ for AZD6738 in *MYC/N* wild-type as compared to *MYC/N* amplified pediatric tumor cell lines (unpaired t-test, *p* > 0.05 for *MYC/N* amp. compared to *MYC/N* amp., not significant, ns). **(D)** Inhibitory concentration 50% values (IC_50_) for AZD6738 in pediatric tumor cell lines expressing high levels of PGBD5 protein (PGBD5 high) as compared to cells expressing low levels of PGBD5 (PGBD5 low) (unpaired t-test, *p* = 0.05 for PGBD5 high compared to PGBD5 low).

**Figure S6.**
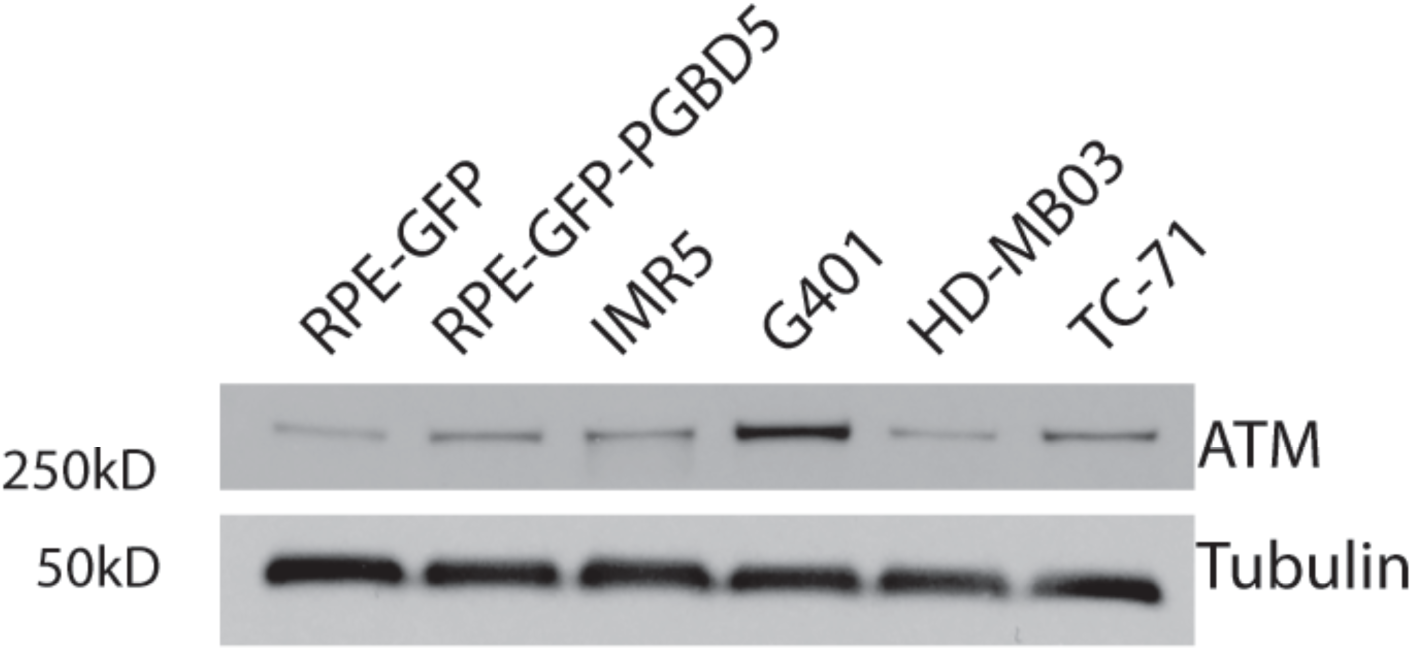
Expression of ATM in PGBD5-expressing normal and tumor cell lines. Tubulin serves as the loading control.

**Figure S7.**
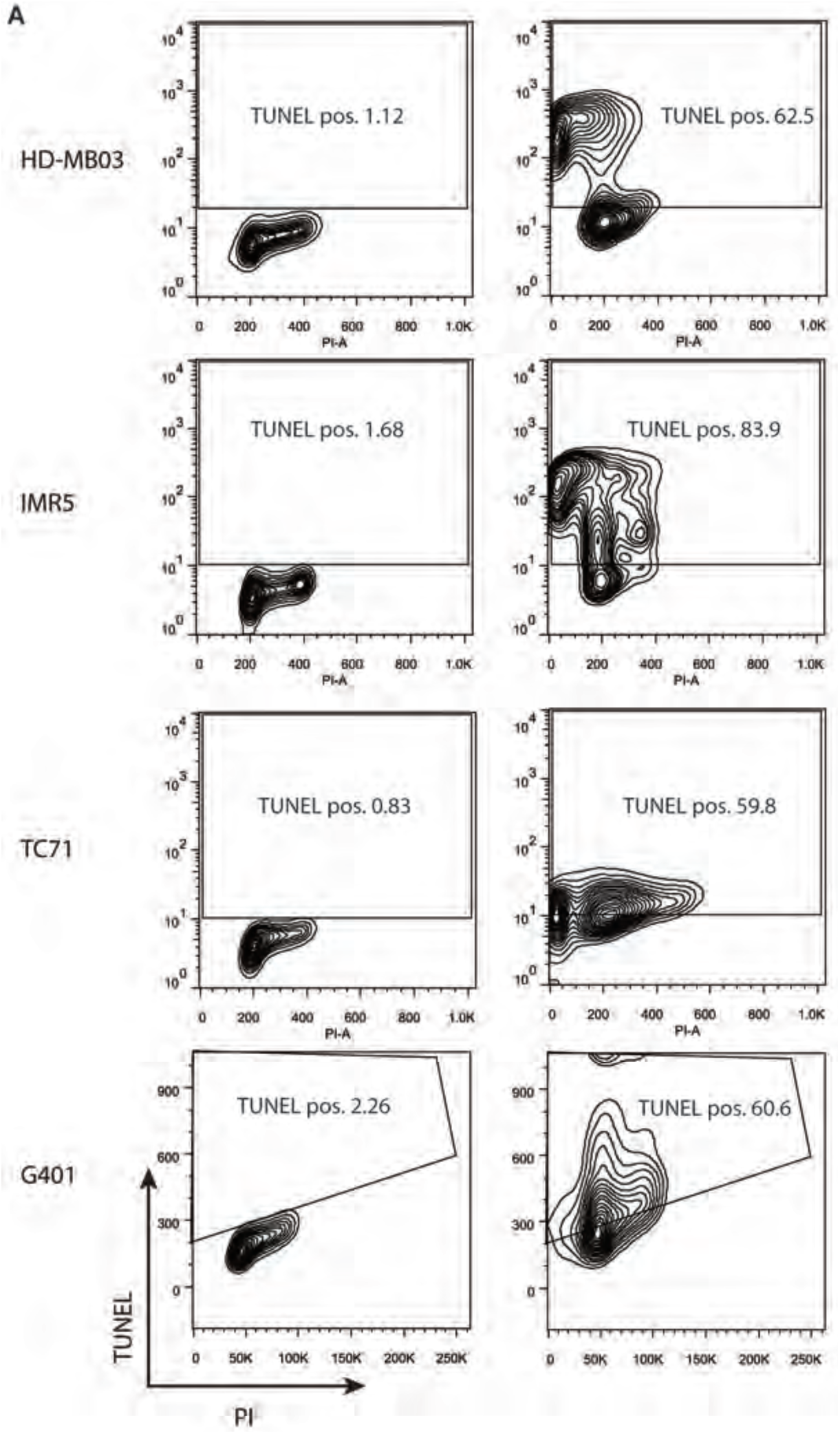
Flow cytometry measurements of cells labeled with TUNEL and treated with AZD6738. Flow cytometric analysis of TUNEL and propidium iodide incorporation into HD-MB03, IMR5, TC71, and G401 cells upon 120 hour treatment of AZD6738 treatment (500 nM).

**Figure S8.**
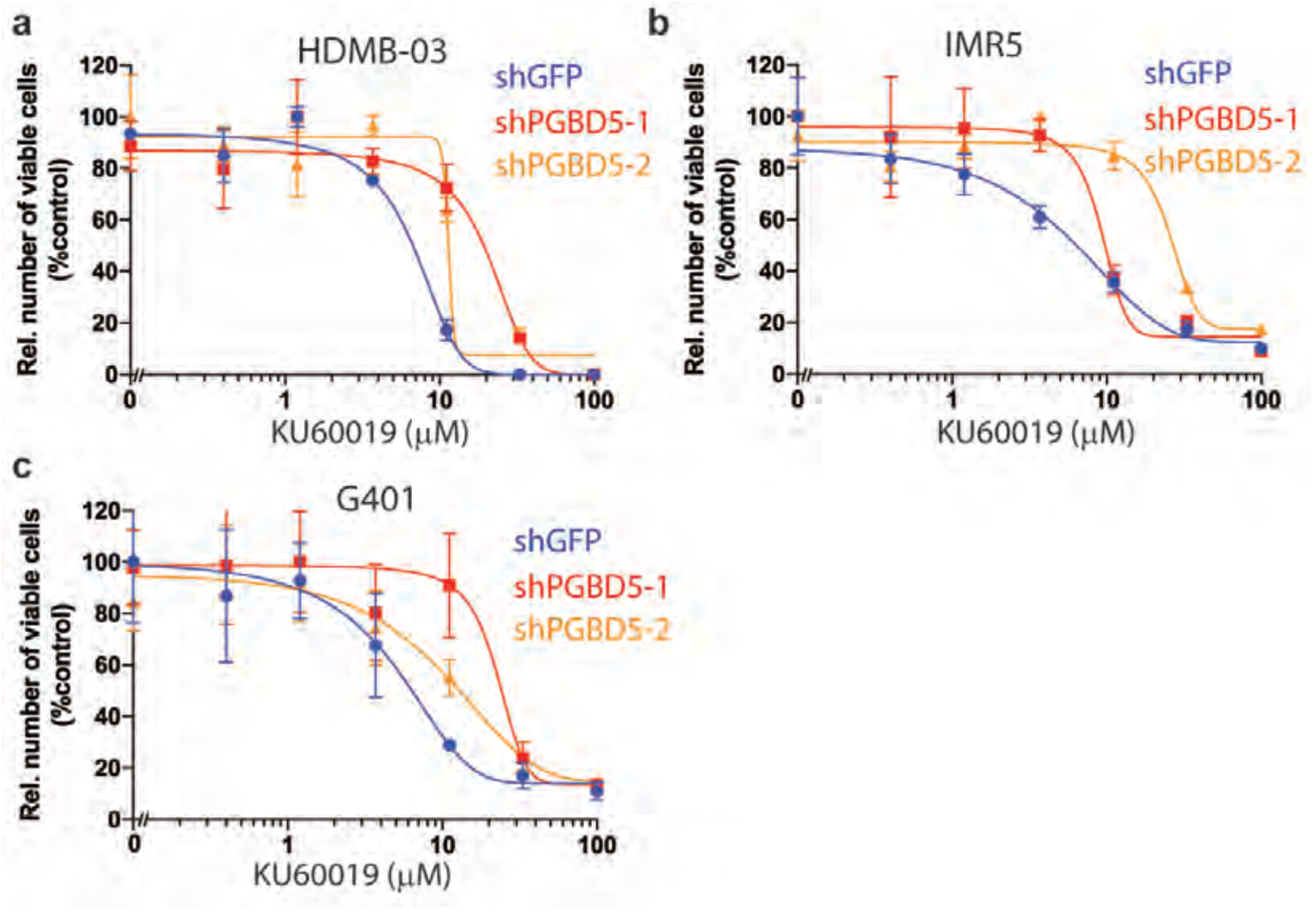
PGBD5 expression is necessary for tumor cell susceptibility to KU60019. (A-C) Dose-response of childhood tumor cell lines HDMB-03 (A), IMR5 (B) and G401 (C) treated with varying concentrations of KU60019 for 120 hours upon *PGBD5* depletion. Error bars represent standard deviations of three replicates.

**Figure S9.**
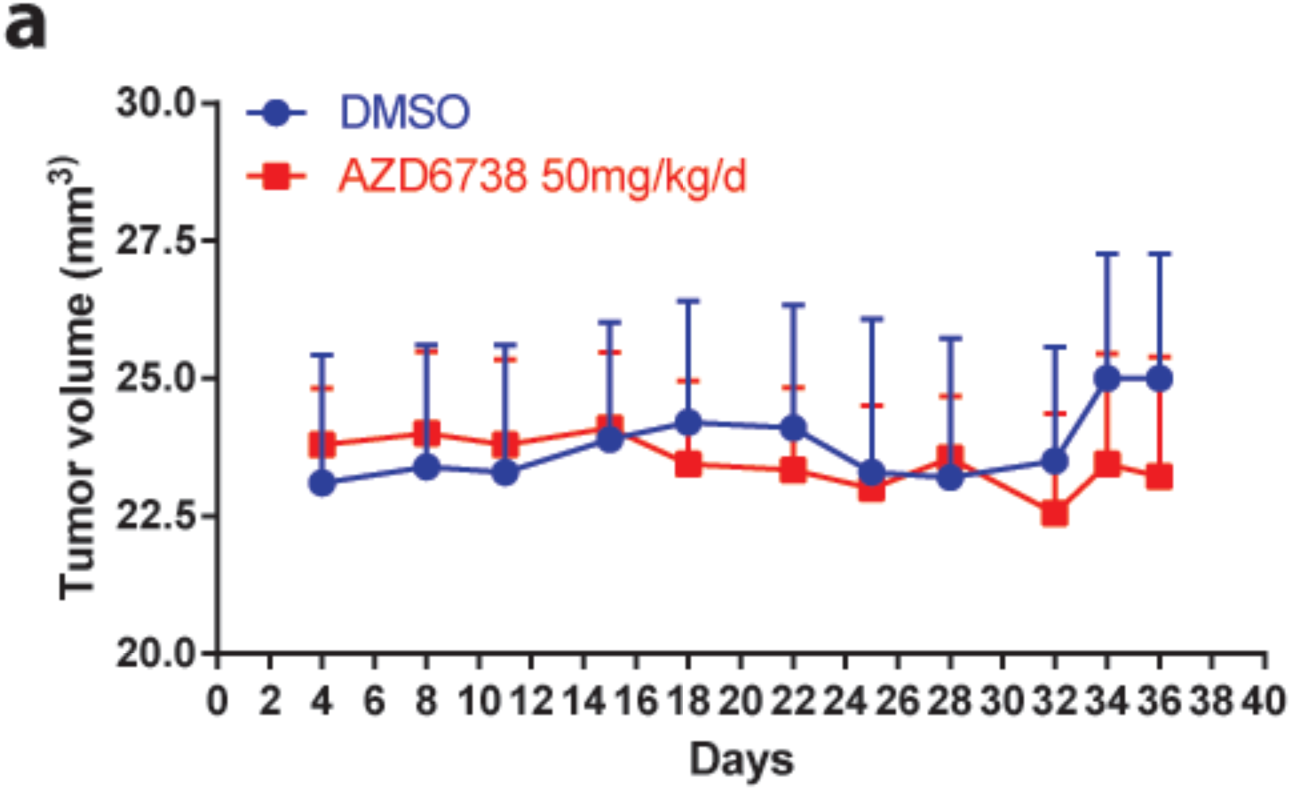
Mouse weight is not affected by treatment with AZD6738. Mouse weights over time in mice treated with AZD6738 at 50 mg/kg daily PO as compared to vehicle control treated mice.

**Figure S10.**
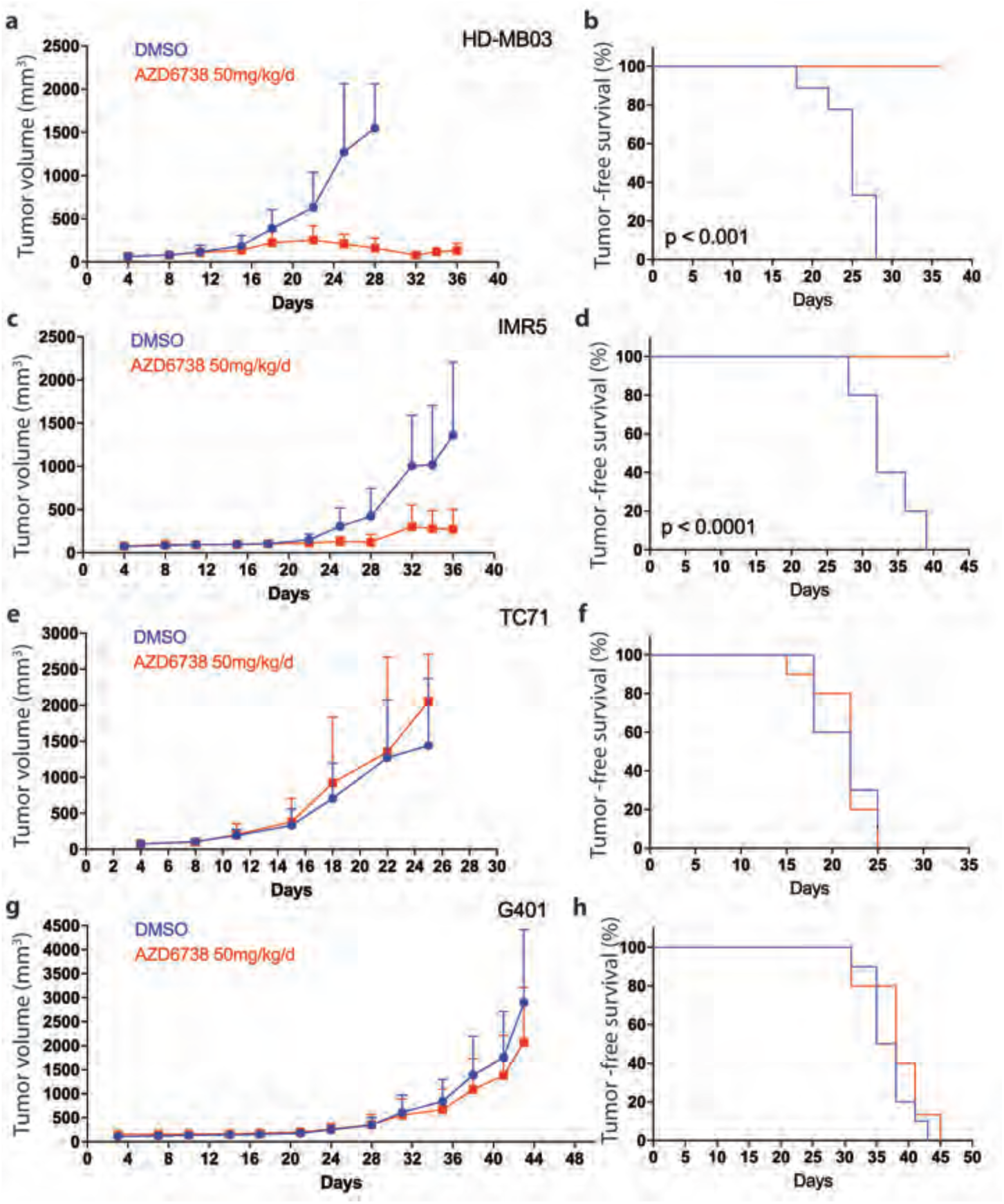
AZD6738 treatment leads to decreased pediatric xenograft tumor burden in mice over time. **(A,C,E,G)** Tumor volumes over time in mice harboring xenografts of HDMB-03 (A), IMR5 (C), TC71 (E) and G401 (G) cells treated with AZD6738 at 50 mg/kg body weight once/day PO as compared to vehicle control treated mice. **(B,D,F,H)** Kaplan Maier analysis of tumor-free survival (defined as tumors <800 mm^3^) in mice harboring xenografts of HDMB-03 (B), IMR5 (D), TC71 (F) and G401 (F) cells treated with AZD6738 at 50 mg/kg body weight once/day PO as compared to vehicle control treated mice.

**Figure S11.**
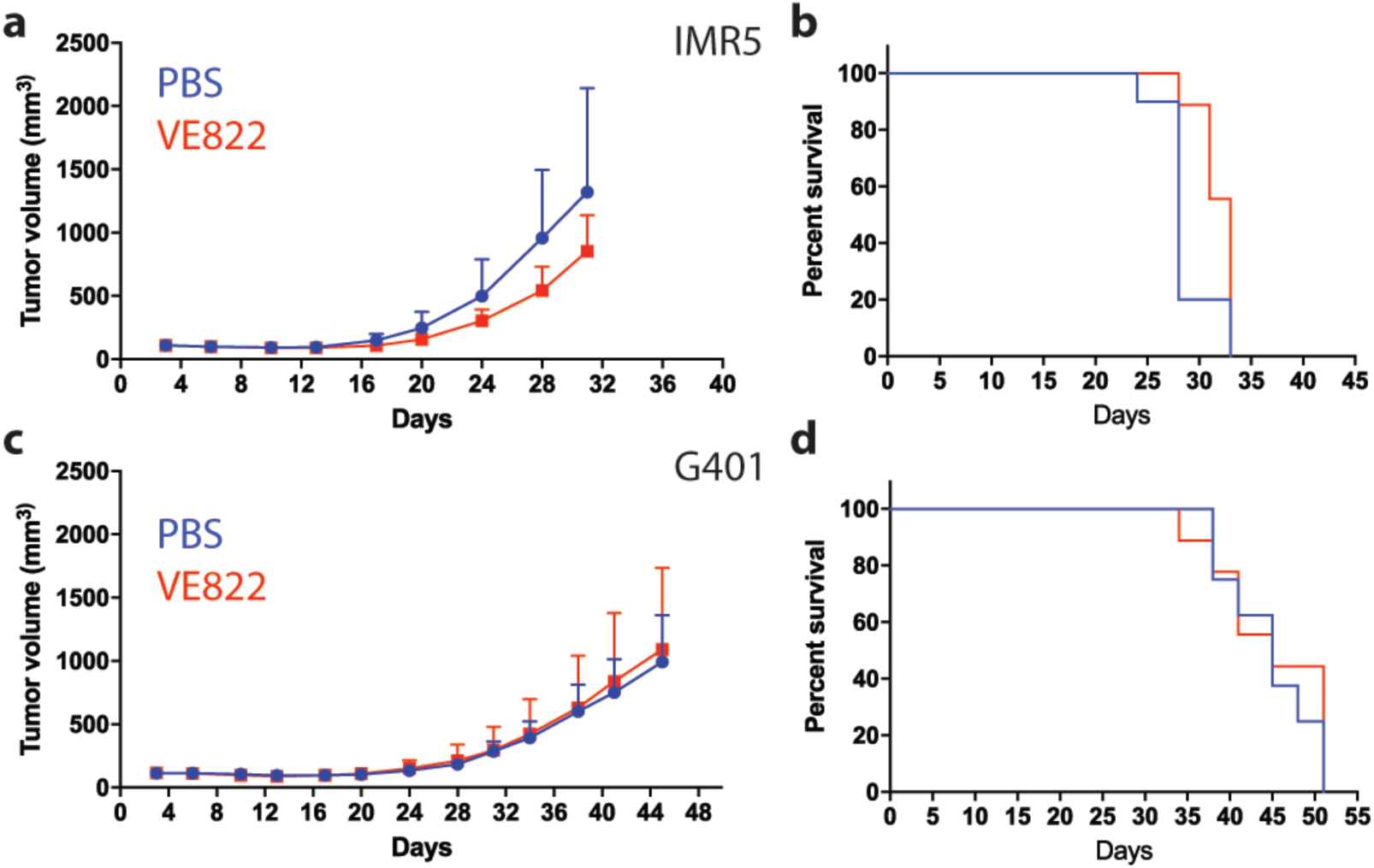
VE822 treatment does not lead to decreased pediatric xenograft tumor burden in mice over time. **(A+C)** Tumor volumes over time in mice harboring xenografts of IMR5 (A) and G401 (C) cells treated with VE822 at 60 mg/kg daily PO (5 days per week), as compared to vehicle control treated mice. **(B+D)** Kaplan Maier analysis of tumor-free survival (defined as tumors <800mm^3^) in mice harboring xenografts of IMR5 (B) and G401 (D) cells treated with VE822 at 60 mg/kg daily PO (5 days per week), as compared to vehicle control treated mice.

**Table S1:** Selectivity profiles of inhibitors of DNA damage signaling.

| Drug | ATM | ATR | DNA-PK | mTOR | Other | Reference |
| --- | --- | --- | --- | --- | --- | --- |
| AZD7762 | N/A | N/A | N/A | N/A | 0.005 (CHK1) | https://doi.org/10.1158/1535-7163.MCT-08-0492 |
| KU60019 | 0.0063 | >10 | 1.7 | <sup>b</sup> | <sup>b,c</sup> | https://doi.org/10.1158/1535-7163.MCT-09-0519 |
| VE822 | 0.034 | <0.0002 | >4 | >1 | 0.22 (PI3Kγ) | https://doi.org/10.1038/cddis.2012.181 |
| AZD6738 | >30<sup>d</sup> | 0.001, 0.074<sup>d</sup> | >30<sup>d</sup> | >23<sup>d</sup> | >30<sup>d</sup> (PI3Kα) | https://doi.org/10.18632/oncotarget.6247 |
| AZ20 | >30<sup>a</sup> | 0.005 | >30<sup>a</sup> | 0.038 | 13 (PI3Kα) | https://doi.org/10.1021/jm301859s |
| NU7441 | >100 | >100 | 0.014 | 2.5 | 5 (PI3Kα) | https://doi.org/10.1016/j.bmcl.2004.09.060 |

Biochemical $K_i$ values are listed in $\mu\text{M}$ , unless otherwise specified. <sup>a</sup> Phenotypic assay against HT29 cells. <sup>b</sup> Supplementary material 2 (Table S1) reports the activity of 1 $\mu\text{M}$ KU60019 as % inhibition, with 104% and 117% activity against mTOR and CHK1, respectively. <sup>c</sup> Almost no activity against PI3K isoforms. <sup>d</sup> Phenotypic assays against cells (Supplementary Table S1).

**Table S2:** Mutational profiles of genes known to affect susceptibility of cells to inhibitors of DNA damage signaling.

|  | IMR5 | CHP-212 | Kelly | SK-N-FI | LAN-1 | HD-MB03 | ONS-76 | Daoy | UW228 | TC-71 | Cado-ES1 | RD-ES1 | G-401 | A-204 | KP-MRT-AN | KP-MRT-RT | KP-MRT-NS | KP-MRT-YM |
| --- | --- | --- | --- | --- | --- | --- | --- | --- | --- | --- | --- | --- | --- | --- | --- | --- | --- | --- |
| ATM | - | - | - | - | - | - | - | - | - | - | - | - | - | - | N/A | N/A | N/A | N/A |
| CHK1 | - | - | - | - | - | - | - | - | - | - | - | - | - | - | N/A | N/A | N/A | N/A |
| MYC/N | mut | - | mut | - | mut | mut | - | - | - | - | - | - | - | - | N/A | N/A | N/A | N/A |
| TP53 | - | - | mut | mut | mut | - | - | - | mut | mut | mut | mut | - | - | N/A | N/A | N/A | N/A |
| BRCA2 | - | - | - | - | - | - | - | - | - | - | - | mut | - | - | N/A | N/A | N/A | N/A |
| XRCC3 | - | - | - | - | - | - | - | - | - | - | - | - | - | - | N/A | N/A | N/A | N/A |
| RAS | - | mut | - | - | mut | - | mut | - | - | - | - | - | - | - | N/A | N/A | N/A | N/A |
| ATR | - | - | - | - | - | - | - | - | - | - | - | - | - | - | N/A | N/A | N/A | N/A |
| ATRX | - | - | - | - | - | - | - | mut | - | - | mut | - | - | - | N/A | N/A | N/A | N/A |
| XRCC5 | - | - | - | - | - | - | - | - | - | - | - | - | - | - | N/A | N/A | N/A | N/A |

Wild-type status is denoted as -. Pathogenic mutations are denoted as mut. Cell lines without mutational data are indicated as N/A.

**Table S3:**
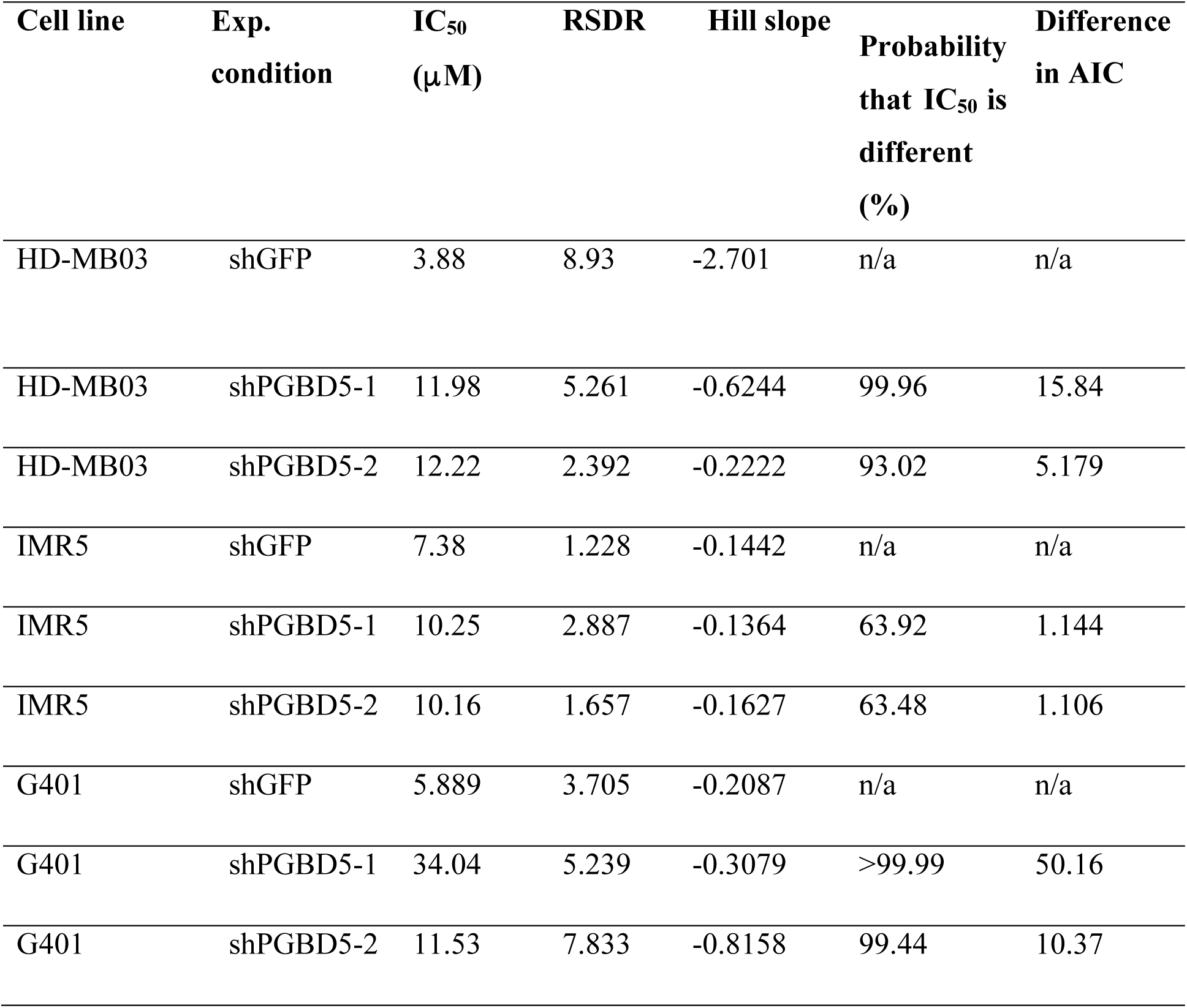
Statistical analysis of drug susceptibility upon PGBD5 depletion.

**Table S4.**
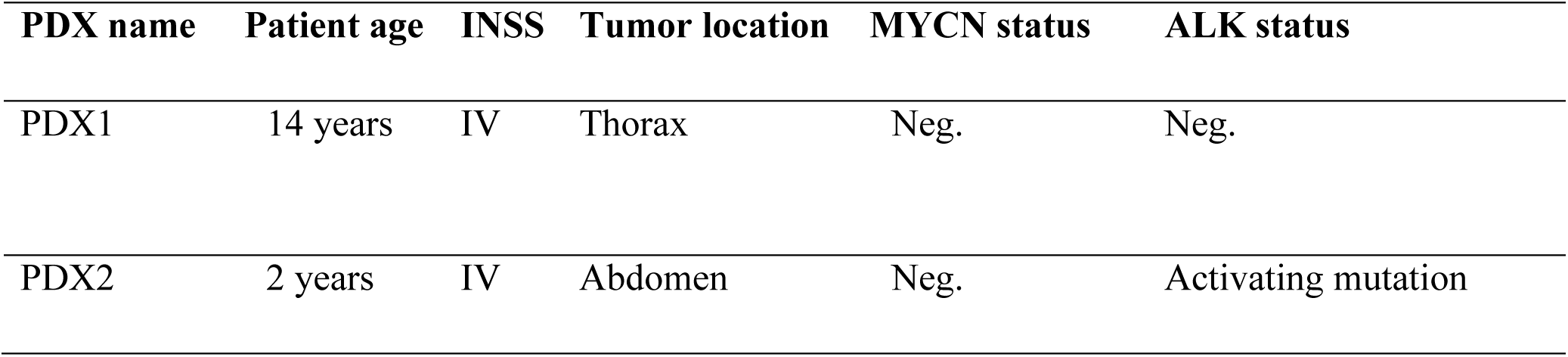
Demographic and molecular features of neuroblastoma patients.

**Table S5:**
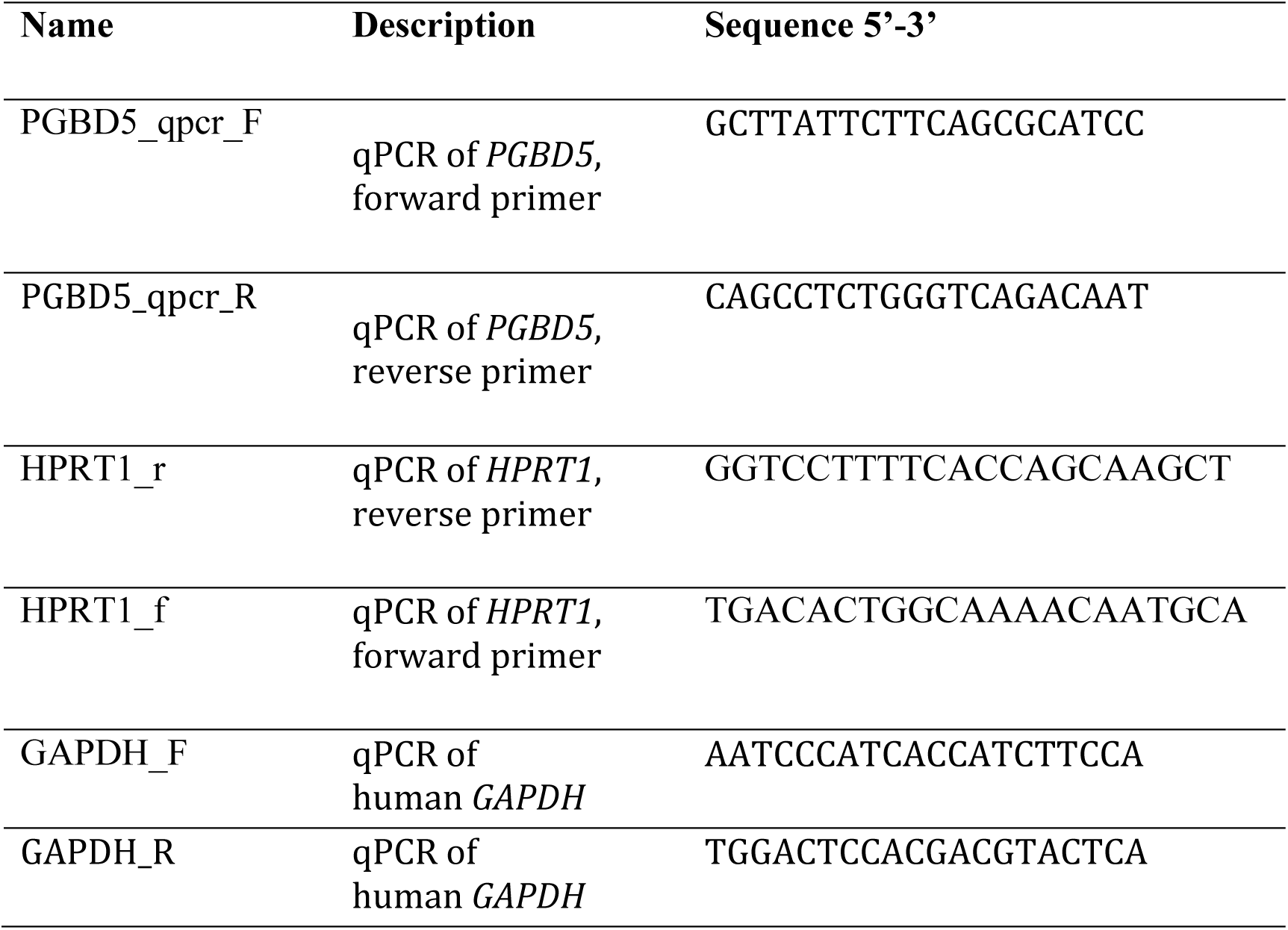
Oligonucleotide primers.

**Table S6:**
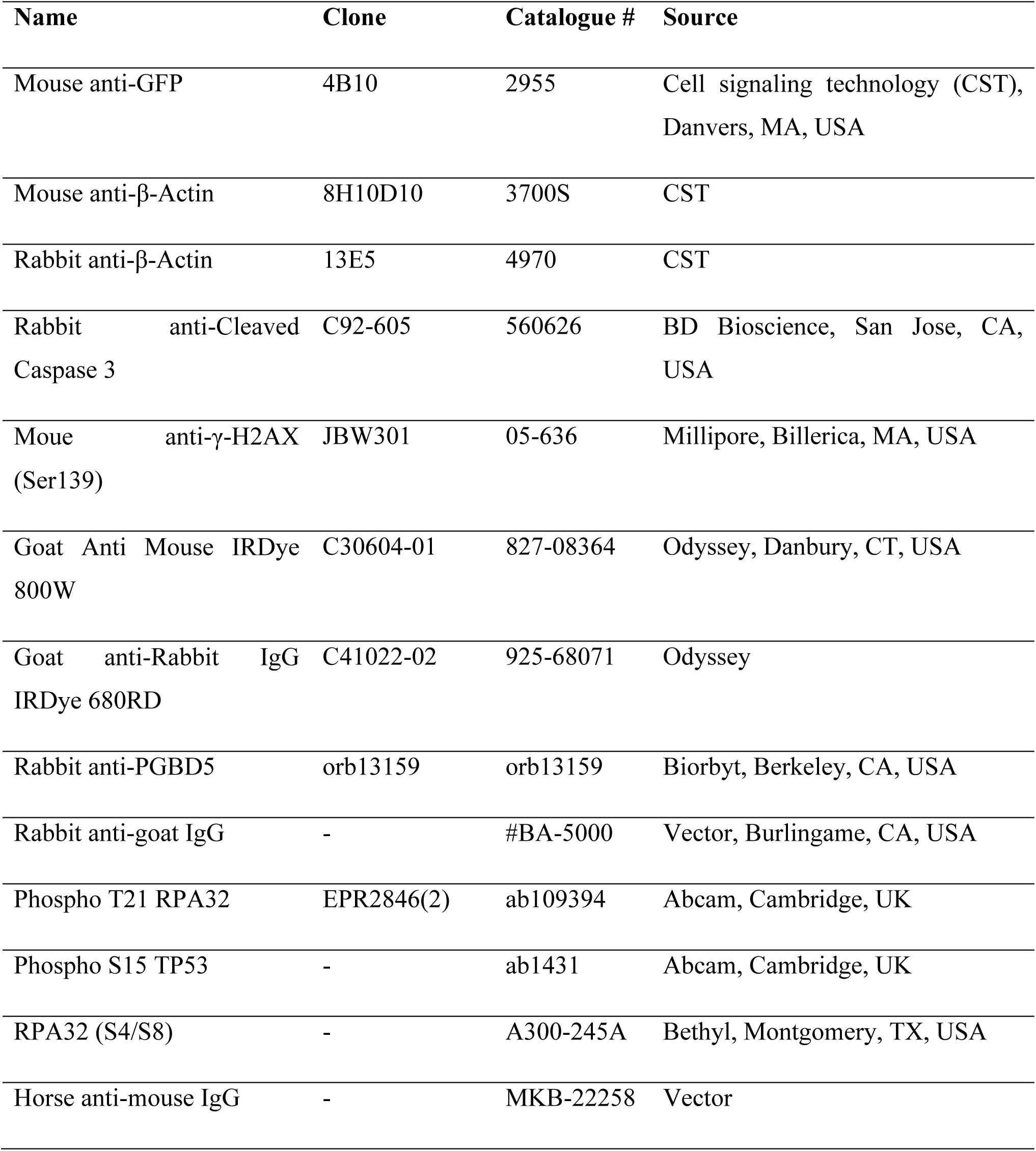
Antibodies.

